# Adipose tissue cell states linked to progression and clinical subtypes of MASLD in human obesity

**DOI:** 10.1101/2025.09.25.678284

**Authors:** J.D. Madsen, T. V. Dam, J. Nambiar, C. Liang, A. R. Peltonen, C.W. Wernberg, L. S. M. Hansen, R. Rydbirk, L. M. Oussoren, K. S. Jakobsen, V. M. G. Larsen, E.G. Klinggaard, L. I. Wassini, F. H. Højgaard, D. Hansen, B. Maniyadath, E. Jonasson, T. D. Caterino, A. Krag, K. Ravnskjaer, S. Mandrup, M. M. Lauridsen, A. Loft, S.F. Schmidt

**Author notes:** These authors contributed equally (J.D. Madsen, T. V. Dam). **Co-correspondence:** Anne Loft, Søren Fisker Schmidt.

## Abstract

Adipose tissue dysfunction is a key determinant of inter-individual variability in obesity-associated comorbidities, yet the underlying tissue-level mechanisms linking adipose remodeling to progression of metabolic dysfunction-associated steatotic liver disease (MASLD) remain poorly defined. We applied single-nucleus RNA sequencing to abdominal subcutaneous adipose tissue (SAT) from individuals with severe obesity, stratified by histologically defined MASLD stage, and integrated these data with bulk RNA-seq from ∼200 individuals. MASLD was associated with adipocyte hypertrophy and near-depletion of an *ADH1B*^HI^ adipocyte subset linked to improved lipid buffering and systemic metabolic health. Conversely, lipid-associated macrophages (LAMs) expanded in MASLD, coordinating lipid-handling and inflammatory programs in the myeloid compartment. In advanced metabolic dysfunction-associated steatohepatitis (MASH), we identified a senescence-associated *PLAU*^HI^ progenitor subpopulation with pro-inflammatory signaling potential. Whereas LAMs were broadly enriched in MASLD subtypes, *PLAU*^HI^ progenitors specifically marked a cardiometabolic MASLD endotype, characterized by type 2 diabetes and cardiovascular disease. These findings highlight cellular remodeling of adipose tissue as a potential driver of MASLD heterogeneity and progression.

## Main

Obesity remains a major global health challenge, fueling a surge in metabolic diseases and their associated complications ^1^. Among these, metabolic dysfunction-associated steatotic liver disease (MASLD) has emerged as the most prevalent liver disorder, affecting approximately 30-40% of the general population and 70% of individuals living with obesity ^2,3^. Nearly half of these cases represent the more severe, inflammatory form of MASLD, metabolic dysfunction-associated steatohepatitis (MASH) ^2^. While MASH rarely causes death directly, its progression to end-stage liver disease and hepatocellular carcinoma contributes to substantial morbidity and mortality. Despite obesity being a major risk factor, the etiological determinants that dictate whether an individual with obesity develops MASH or remains in the relatively benign state of hepatic steatosis are still poorly understood. Adding to this complexity is the clinical heterogeneity of MASH, which has recently been stratified into distinct endotypes ^4^. Emerging evidence points to several key drivers of disease progression, including genetic predisposition, insulin resistance, lipotoxicity, and organ crosstalk, particularly between adipose tissue and the liver ^5^.

Adipose tissue serves as a major energy reservoir and plays a central role in maintaining systemic energy homeostasis ^6^. In metabolically healthy individuals, subcutaneous adipose tissue (SAT) normally functions as the primary long-term lipid storage depot, buffering systemic energy excess, which is associated with metabolic flexibility and protection against insulin resistance ^7,8^. When SAT becomes dysfunctional, characterized by impaired lipid storage, impaired adipogenesis, and altered secretory profiles, excess lipids may spill over to ectopic sites, promoting hepatic lipid overload and systemic metabolic dysfunction ^6^. Evidence from mouse models and patients with lipodystrophy shows that loss of healthy adipose tissue function promotes metabolic disease even in the absence of obesity ^9–11^, whereas preserved SAT function can mitigate complications despite excess adiposity ^12^. While dysfunctional SAT is recognized as a central driver of ectopic lipid accumulation in the liver in MASLD, its role in shaping the transition from hepatic steatosis to MASH is not well understood. Furthermore, although previous studies using bulk RNA-seq ^13^ have provided insights into changes in the SAT transcriptome across MASLD stages ^14,15^, a comprehensive, cell type–resolved dissection of SAT remodeling at the whole-tissue level has not yet been achieved.

Here, we applied single-nucleus RNA sequencing (snRNA-seq) to abdominal SAT from individuals with severe obesity, stratified by liver histology status. We identified cellular subpopulations, states and gene programs in SAT associated with MASLD severity and clinical MASH endotypes, providing new cell type-resolved insights into the link between adipose tissue function and MASLD severity in human obesity. These findings suggest that adipose tissue dysfunction may drive MASLD progression and highlight the potential of targeting specific cell states to preserve metabolic function and mitigate fatty liver disease.

## Results

### Compositional remodeling of cell types in SAT with increasing MASLD severity

To investigate the heterogeneity of SAT in individuals with obesity, we collected human abdominal SAT and liver biopsies from a cohort of 67 female patients with severe obesity ^16^ with a median body mass index (BMI) of 42. Based on histological assessment of the liver biopsies obtained utilizing the Steatosis-Activity-Fibrosis (SAF) score ^17^, individuals with obesity were stratified into three groups; individuals with normal liver histology (SAF1, no-MASLD; n = 22), individuals with steatosis (SAF2, steatosis; n = 25), and individuals with MASH (SAF3, MASH; n = 20) (Figure 1A-B). There were no differences in BMI, age or body fat percentage between these patient groups, whereas liver fat percentage, NAS score and liver fibrosis (as assessed by liver elastography) were elevated in patient groups with steatosis and MASH compared to the group of patients without MASLD (Table 1). To assess the transcriptional differences between the patient groups, we performed bulk RNA-sequencing of the SAT. This revealed substantial transcriptional differences between the disease stages (padj < 0.1) linked to higher expression of genes involved in inflammatory pathways and tissue fibrosis and lower expression of genes coupled to metabolic pathways in SAT with increasing SAF score (Figure 1C-D). While clear transcriptional differences existed between patient groups, considerable heterogeneity was also evident within each group, which was especially evident for the cluster containing inflammatory genes (Figure 1C). To identify representative SAT RNA-seq profiles for each SAF diagnosis, we conducted linear discriminant analysis (LDA) on the top 500 most variable genes in the RNA-seq data, followed by archetype analysis (Figure 1E and S1A). When selecting three archetypes, most patients were grouped in closest proximity to their SAF-enriched archetype, although a few patients within each SAF score diagnosis appeared to be better or equally well described by an alternative archetype (Figure 1E). Subsequently, we selected eight patients from each group with the highest corresponding archetype coefficients for single-cell analyses (Figure S1B). Importantly, there was a strong concordance in both the differentially expressed genes and the clinical parameters between patient groups in SAT for the selected archetypical females and the entire cohort of female patients (Figure 1F-H and S1C-F), suggesting that the 24 archetypical females are reliable representatives of the whole patient cohort. To minimize batch effects during preparation of SAT samples for snRNA-seq, we pooled female donors ensuring equal representation from each patient group prior to snRNA-seq analyses and used a genetic demultiplexing strategy to bioinformatically assign nuclei to each donor after sequencing (see Methods). Following strict quality filtering steps, we obtained 36.846 nuclei, recovering all major cell types in SAT with a strong donor representation in each cell cluster (Figure 1I and S1G-H). The cell clusters were defined by a distinct set of established markers ^18^, such as *PLIN1* for adipocytes, *MRC1* for myeloid cells, *PDGFRA* for adipose stromal and progenitor cells (ASPCs), and *MECOM* for endothelial cells, which could be linked to known functional properties of each cell type (Figure S1I and Table S1-2). We next evaluated the compositional differences in SAT in patient groups stratified by SAF score. Interestingly, the proportion of adipocytes trended to be lower in patients with steatosis and MASH compared to patients without MASLD, while the proportion of myeloid cells trended to be higher in both patients with steatosis and MASH (Figure 1J). To further substantiate these observations, we applied MiloR ^19^, which determines differences in cell type abundance across experimental conditions by analyzing neighborhoods of cells in the k-nearest neighbor graph. Consistent with our initial findings, many myeloid cell-associated neighborhoods show increased abundance in patients with steatosis and MASH compared to individuals without MASLD (Figure S1J). In contrast, many adipocyte-related neighborhoods exhibit reduced abundance in steatosis and MASH, aligning with the overall decrease in adipocyte fraction. However, a subset of these neighborhoods shows significantly increased abundance, indicating a heterogeneous response of the adipocyte compartment in SAT during MASLD progression in individuals with obesity. To investigate compositional changes in the larger cohort, we applied a deconvolution algorithm ^20^ to the bulk RNA-seq data from all 67 obese female patients with well-defined MASLD states, confirming an overall reduction in adipocyte fractions and a trending increase in myeloid cell fractions in patients with steatosis and MASH compared to individuals without MASLD (Figure 1K). To corroborate our computational findings, we performed a histological staining of SAT to assess adipocyte size and number per area across varying SAF scores. Here we found that adipocyte size was increased in patients with steatosis and MASH (Figure 1L-M and S1K) and correlates with increased liver fat observed in these two patient groups (Figure 1N). Collectively, this indicates that the overall decrease in adipocyte fraction is paralleled by a hypertrophic expansion of adipocytes as MASLD progresses.

**Figure 1.**
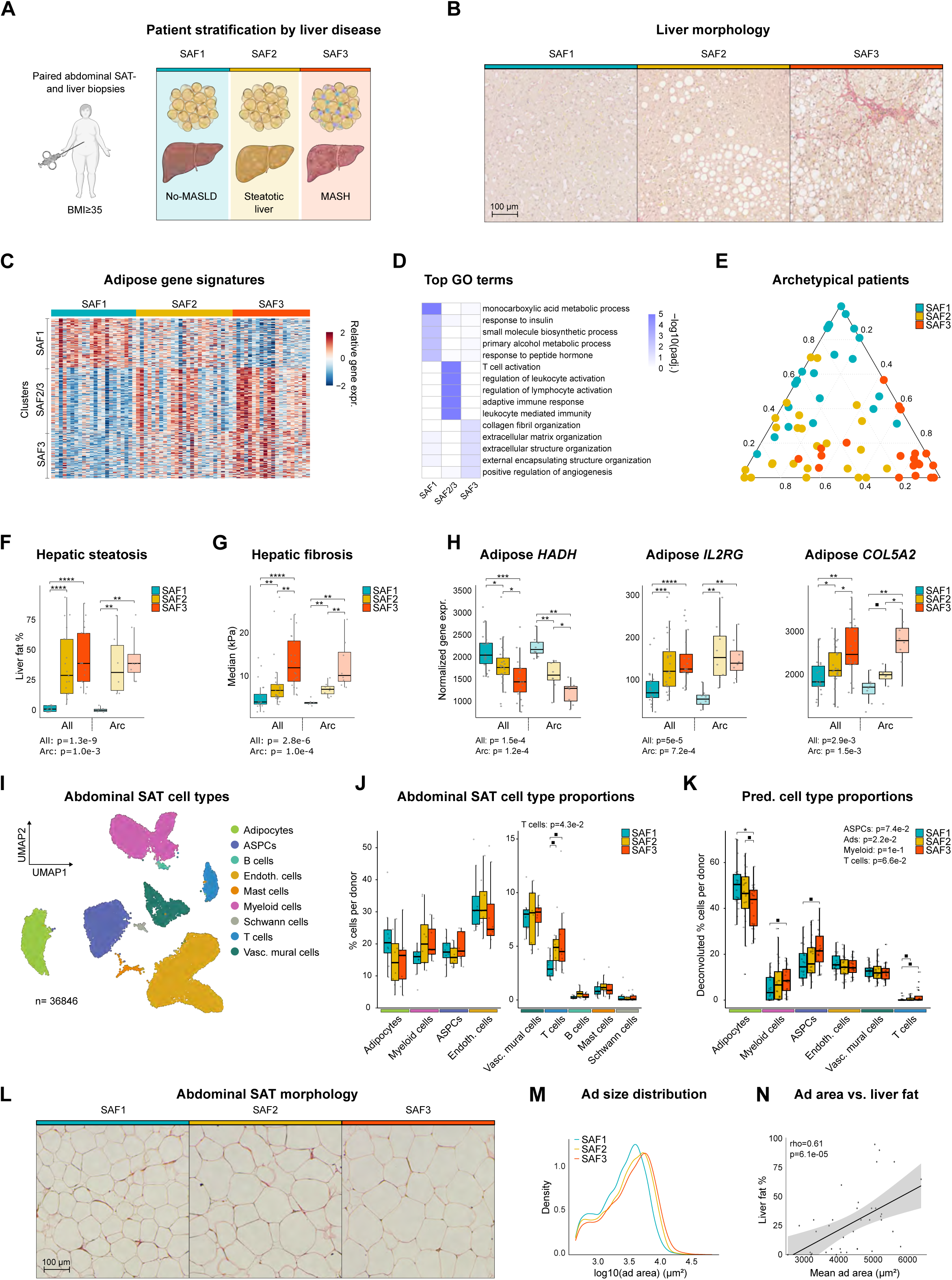
| Transcriptomic and histological profiling of females with obesity stratified by MASLD severity. **A**, Abdominal subcutaneous adipose tissue (SAT) and liver biopsies were obtained from 67 females with severe obesity (BMI > 35) and stratified into no-MASLD (SAF1), steatosis (SAF2), and MASH (SAF3) groups based on biopsy-proven Steatosis-Activity-Fibrosis (SAF) scores. **B**, Representative Sirius Red-stained liver sections from each SAF category. Scale bars, 100 μm. **C**, K-means clustering (k = 3) of row-scaled normalized expression counts of genes differentially expressed (padj < 0.1) between one or more SAF groups. **D**, Gene ontology (GO) biological process enrichment analysis of RNA-seq clusters shown in (**C**). **E**, Ternary plot showing archetype scores for each individual female patient, colored by SAF diagnosis. **F–G**, Quantification of liver fat percentage (all: S1, n=21; S2, n=21, S3, n=18; archetype, S1, n=7; S2, n=6, S3, n=8) (**F**) and hepatic fibrosis signal (all: S1, n=21; S2, n=25, S3, n=20; archetype, n=8 per condition) (**G**). **H**, Normalized expression of representative genes from each RNA-seq cluster (all: S1, n=22; S2, n=25, S3, n=20; archetype, n=8 per condition). **I**, UMAP visualization of 36,846 SAT nuclei from all donors. **J**, Proportion of a specific cell population relative to all cells in the female SAT snRNA-seq dataset (n=8 per condition). **K**, Deconvoluted cell type proportions in the female SAT bulk RNA-seq dataset (S1, n=22; S2, n=25, S3, n=20). **L**, Representative Masson-Trichrome-stained SAT sections. Scale bars, 100 μm. **M**, Distribution of adipocyte area. **N,** Correlation between mean adipocyte area and liver fat percentage. Boxplot data: center line, median; box limits, upper and lower quartiles; whiskers, 1.5x IQR. A Kruskal-Wallis (KW) test, followed by a post-hoc Wilcoxon rank-sum test with Benjamini-Hochberg (BH) correction for multiple testing, was performed to assess significance between SAF diagnosis for **F-H** and **J-K** (▪, p<0.1; *, p<0.05; **, p<0.01; ***, p<0.001; ****, p<0.0001). Correlation analyses were performed using Spearman correlation; reported p-values are based on two-sided tests in **N**. ASPCs, adipogenic stromal and progenitor cells. Credit: a, Created with BioRender.com.

**Table 1.**
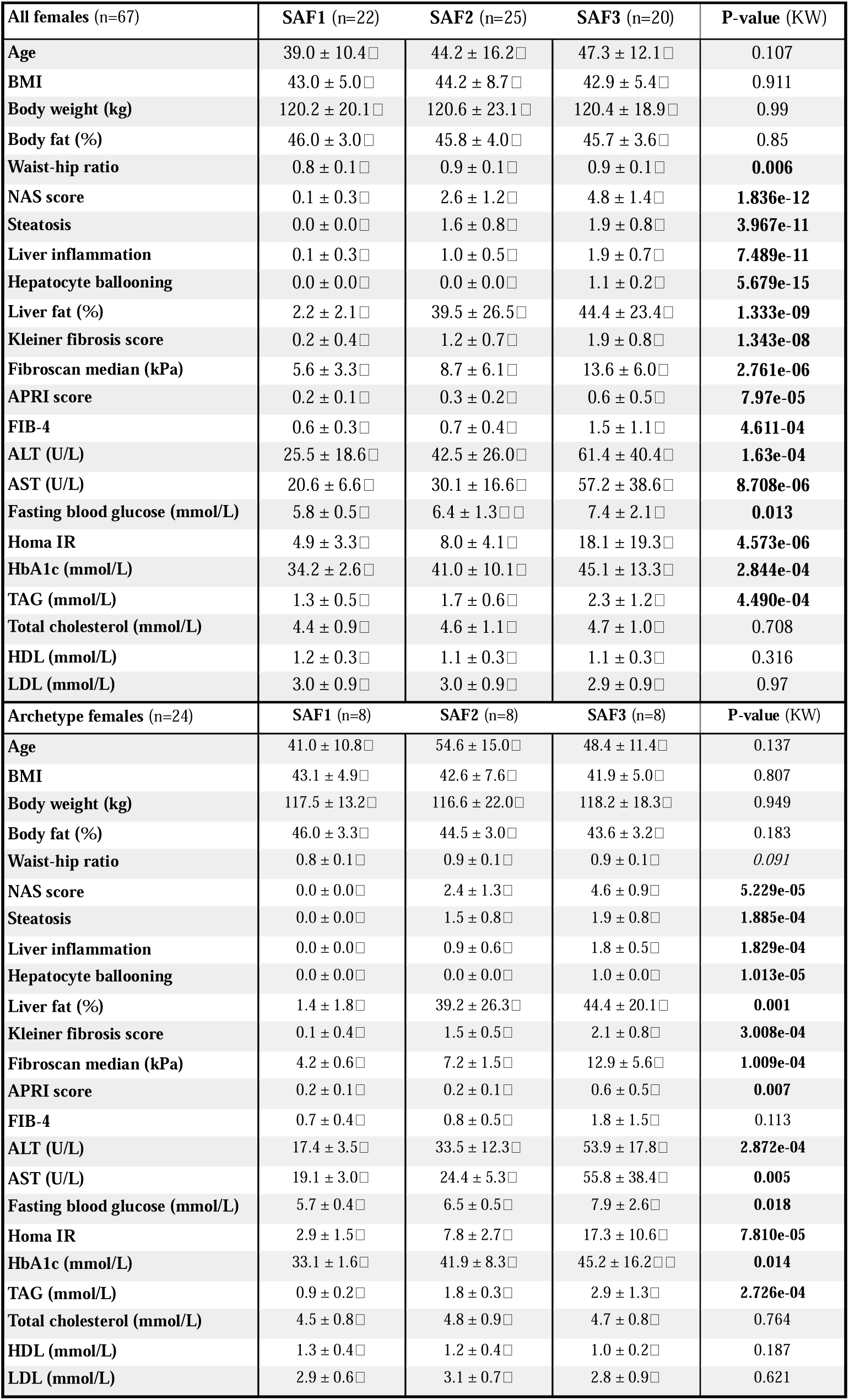
| Clinical, biochemical, and functional characteristics of patients. The table indicates mean values (±SD) for 67 females (upper panel) and 24 archetypical females (lower panel) stratified by SAF score. A Kruskal - Wallis (KW) test, followed by a posthoc Wilcoxon rank-sum test with Benjamin Hochberg (BH) correction for multiple testing, was performed to assess significance between SAF diagnosis. Different letters indicate significant differences between groups (padj < 0.05), whereas shared letters denote no significant difference. ALT, alanine aminotransferase; APRI, aspartate aminotransferase to platelet ratio index; AST, aspartate aminotransferase; BMI, body mass index; FIB-4, fibrosis-4 index; HbA1c, glycated hemoglobin A1c; HDL, high-density lipoprotein; HOMA-IR, homeostatic model assessment of insulin resistance; LDL, low-density lipoprotein; NAS, nonalcoholic fatty liver disease activity score; SAF, steatosis, activity, and fibrosis score; TAG, triacylglycerol.

| All females (n=67) | SAF1 (n=22) | SAF2 (n=25) | SAF3 (n=20) | P-value (KW) |
| --- | --- | --- | --- | --- |
| Age | 39.0 ± 10.4□ | 44.2 ± 16.2□ | 47.3 ± 12.1□ | 0.107 |
| BMI | 43.0 ± 5.0□ | 44.2 ± 8.7□ | 42.9 ± 5.4□ | 0.911 |
| Body weight (kg) | 120.2 ± 20.1□ | 120.6 ± 23.1□ | 120.4 ± 18.9□ | 0.99 |
| Body fat (%) | 46.0 ± 3.0□ | 45.8 ± 4.0□ | 45.7 ± 3.6□ | 0.85 |
| Waist-hip ratio | 0.8 ± 0.1□ | 0.9 ± 0.1□ | 0.9 ± 0.1□ | **0.006** |
| NAS score | 0.1 ± 0.3□ | 2.6 ± 1.2□ | 4.8 ± 1.4□ | **1.836e-12** |
| Steatosis | 0.0 ± 0.0□ | 1.6 ± 0.8□ | 1.9 ± 0.8□ | **3.967e-11** |
| Liver inflammation | 0.1 ± 0.3□ | 1.0 ± 0.5□ | 1.9 ± 0.7□ | **7.489e-11** |
| Hepatocyte ballooning | 0.0 ± 0.0□ | 0.0 ± 0.0□ | 1.1 ± 0.2□ | **5.679e-15** |
| Liver fat (%) | 2.2 ± 2.1□ | 39.5 ± 26.5□ | 44.4 ± 23.4□ | **1.333e-09** |
| Kleiner fibrosis score | 0.2 ± 0.4□ | 1.2 ± 0.7□ | 1.9 ± 0.8□ | **1.343e-08** |
| Fibroscan median (kPa) | 5.6 ± 3.3□ | 8.7 ± 6.1□ | 13.6 ± 6.0□ | **2.761e-06** |
| APRI score | 0.2 ± 0.1□ | 0.3 ± 0.2□ | 0.6 ± 0.5□ | **7.97e-05** |
| FIB-4 | 0.6 ± 0.3□ | 0.7 ± 0.4□ | 1.5 ± 1.1□ | **4.611-04** |
| ALT (U/L) | 25.5 ± 18.6□ | 42.5 ± 26.0□ | 61.4 ± 40.4□ | **1.63e-04** |
| AST (U/L) | 20.6 ± 6.6□ | 30.1 ± 16.6□ | 57.2 ± 38.6□ | **8.708e-06** |
| Fasting blood glucose (mmol/L) | 5.8 ± 0.5□ | 6.4 ± 1.3□□ | 7.4 ± 2.1□ | **0.013** |
| Homa IR | 4.9 ± 3.3□ | 8.0 ± 4.1□ | 18.1 ± 19.3□ | **4.573e-06** |
| HbA1c (mmol/L) | 34.2 ± 2.6□ | 41.0 ± 10.1□ | 45.1 ± 13.3□ | **2.844e-04** |
| TAG (mmol/L) | 1.3 ± 0.5□ | 1.7 ± 0.6□ | 2.3 ± 1.2□ | **4.490e-04** |
| Total cholesterol (mmol/L) | 4.4 ± 0.9□ | 4.6 ± 1.1□ | 4.7 ± 1.0□ | 0.708 |
| HDL (mmol/L) | 1.2 ± 0.3□ | 1.1 ± 0.3□ | 1.1 ± 0.3□ | 0.316 |
| LDL (mmol/L) | 3.0 ± 0.9□ | 3.0 ± 0.9□ | 2.9 ± 0.9□ | 0.97 |
| Archetype females (n=24) | SAF1 (n=8) | SAF2 (n=8) | SAF3 (n=8) | P-value (KW) |
| Age | 41.0 ± 10.8□ | 54.6 ± 15.0□ | 48.4 ± 11.4□ | 0.137 |
| BMI | 43.1 ± 4.9□ | 42.6 ± 7.6□ | 41.9 ± 5.0□ | 0.807 |
| Body weight (kg) | 117.5 ± 13.2□ | 116.6 ± 22.0□ | 118.2 ± 18.3□ | 0.949 |
| Body fat (%) | 46.0 ± 3.3□ | 44.5 ± 3.0□ | 43.6 ± 3.2□ | 0.183 |
| Waist-hip ratio | 0.8 ± 0.1□ | 0.9 ± 0.1□ | 0.9 ± 0.1□ | *0.091* |
| NAS score | 0.0 ± 0.0□ | 2.4 ± 1.3□ | 4.6 ± 0.9□ | **5.229e-05** |
| Steatosis | 0.0 ± 0.0□ | 1.5 ± 0.8□ | 1.9 ± 0.8□ | **1.885e-04** |
| Liver inflammation | 0.0 ± 0.0□ | 0.9 ± 0.6□ | 1.8 ± 0.5□ | **1.829e-04** |
| Hepatocyte ballooning | 0.0 ± 0.0□ | 0.0 ± 0.0□ | 1.0 ± 0.0□ | **1.013e-05** |
| Liver fat (%) | 1.4 ± 1.8□ | 39.2 ± 26.3□ | 44.4 ± 20.1□ | **0.001** |
| Kleiner fibrosis score | 0.1 ± 0.4□ | 1.5 ± 0.5□ | 2.1 ± 0.8□ | **3.008e-04** |
| Fibroscan median (kPa) | 4.2 ± 0.6□ | 7.2 ± 1.5□ | 12.9 ± 5.6□ | **1.009e-04** |
| APRI score | 0.2 ± 0.1□ | 0.2 ± 0.1□ | 0.6 ± 0.5□ | **0.007** |
| FIB-4 | 0.7 ± 0.4□ | 0.8 ± 0.5□ | 1.8 ± 1.5□ | 0.113 |
| ALT (U/L) | 17.4 ± 3.5□ | 33.5 ± 12.3□ | 53.9 ± 17.8□ | **2.872e-04** |
| AST (U/L) | 19.1 ± 3.0□ | 24.4 ± 5.3□ | 55.8 ± 38.4□ | **0.005** |
| Fasting blood glucose (mmol/L) | 5.7 ± 0.4□ | 6.5 ± 0.5□ | 7.9 ± 2.6□ | **0.018** |
| Homa IR | 2.9 ± 1.5□ | 7.8 ± 2.7□ | 17.3 ± 10.6□ | **7.810e-05** |
| HbA1c (mmol/L) | 33.1 ± 1.6□ | 41.9 ± 8.3□ | 45.2 ± 16.2□□ | **0.014** |
| TAG (mmol/L) | 0.9 ± 0.2□ | 1.8 ± 0.3□ | 2.9 ± 1.3□ | **2.726e-04** |
| Total cholesterol (mmol/L) | 4.5 ± 0.8□ | 4.8 ± 0.9□ | 4.7 ± 0.8□ | 0.764 |
| HDL (mmol/L) | 1.3 ± 0.4□ | 1.2 ± 0.4□ | 1.0 ± 0.2□ | 0.187 |
| LDL (mmol/L) | 2.9 ± 0.6□ | 3.1 ± 0.7□ | 2.8 ± 0.9□ | 0.621 |

### Cell type-specific transcriptional rewiring of SAT during MASLD development

While prior studies have examined transcriptional changes in human adipose tissue across the MASLD spectrum using bulk analyses ^14,15,21,22^ or focused on macrophages ^13^ in visceral fat depots, single cell-resolved understanding of transcriptional SAT remodeling in obesity with and without MASLD has not been achieved. Using the snRNA-seq data, we examined transcriptional differences across major SAT cell types in MASLD by performing pseudobulk analysis within each population. The most prominent transcriptional differences between the distinct disease stages were observed in myeloid cells, adipose stromal and progenitor cells (ASPCs), and adipocytes (Figure S2A). Despite a similar number of endothelial cells in our dataset, only a few significant transcriptional changes were observed in this population, suggesting that endothelial cells do not undergo significant transcriptional remodeling in response to MASLD progression (Figure S2A).

We next clustered genes differentially expressed between SAF1, SAF2, and/or SAF3 within each cell type (padj < 0.1) and identified two major MASLD-regulated clusters in ASPCs and three in both adipocytes and myeloid cells (Figure 2A-C and Table S3). While most genes and gene programs exhibited cell type-specific regulation across clusters (Figure 2D–G and S2B–F), MASH-enriched genes showed a higher degree of overlap particularly between ASPCs and adipocytes (Figure S2D). Shared MASH-enriched gene programs are linked to inflammatory responses, and the shared genes encode transcriptional regulators such as RFX2, FOSL2, BCL3, and RELB (Figure S2E), which have recently been implicated in an obesity-induced stress response in human SAT ^23^. Clusters with elevated expression in patients without MASLD (i.e., a ‘SAF1 signature’) are enriched for metabolic pathways in adipocytes and G protein-coupled receptor signaling and cell migration in myeloid cells (Figure 2E-F), suggesting that core functional programs in these cell types are progressively silenced in SAT as MASLD progresses. In ASPCs, most repressed genes are not downregulated until the MASH stage; however, some genes, such as *EBF2*, a key transcriptional regulator in progenitor cells ^24^, are already markedly suppressed in individuals with steatosis (Figure 2F), suggesting early disruption of adipogenic programming in MASLD. The cluster with high expression in patients with steatosis and MASH (i.e., a ‘SAF2/3 signature’) in adipocytes is associated with extracellular matrix organization and includes multiple collagen-encoding genes (Figure 2E and G). In myeloid cells, the SAF2/3-associated cluster is enriched for lipid and cholesterol storage pathways (Figure 2E and G), suggesting an increased capacity for lipid handling in this cell type, with advancing disease severity paralleling the observed adipocyte hypertrophy (Figure 1L-M). Taken together, MASLD severity is associated with comprehensive cell type-specific transcriptional reprogramming in SAT, primarily in myeloid cells, ASPCs and adipocytes.

**Figure 2.**
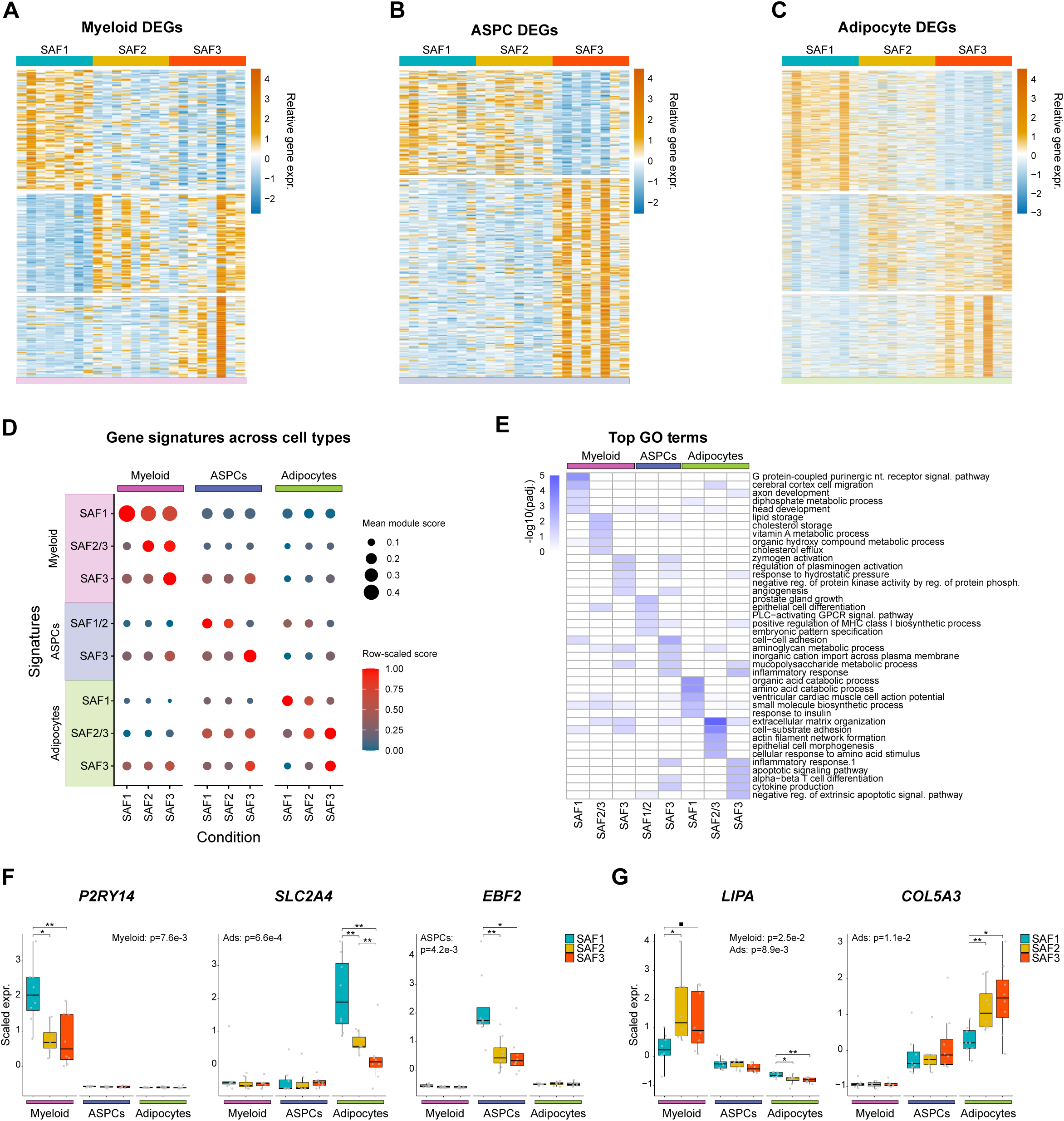
| Cell type-specific transcriptomic signatures in SAT associated with MASLD severity. **A–C**, K-means clustering of normalized pseudobulk expression of differentially expressed genes (DEGs) identified in myeloid immune cells (**A**), adipose stromal and progenitor cells (ASPCs; **B**), and adipocytes (**C**) across SAF categories. **D**, Gene module score of gene clusters identified in (**A**–**C**) across cell types and conditions. Dot size reflects mean module score; color indicates row-scaled gene module score. **E**, Gene ontology (GO) biological process enrichment analysis showing the top five terms for each RNA-seq cluster from (**A**-**C**). **F–G**, Normalized pseudobulk expression of representative cell type-specific cluster genes with SAF1-(**F**) and SAF2/3-(**G**) associated signatures (n=8 per condition). Boxplot data: center line, median; box limits, upper and lower quartiles; whiskers, 1.5x IQR. A KW test, followed by a post-hoc Wilcoxon rank-sum test with BH correction for multiple testing, was performed to assess significance between SAF diagnosis for **F-G** (▪, p<0.1; *, p<0.05; **, p<0.01). ASPCs, adipogenic stromal and progenitor cells.

### Lipid-associated macrophages are elevated in patients with hepatic steatosis and MASH

Inflammatory cells in adipose tissue, particularly certain types of macrophages, are strongly linked to BMI ^23,25^, highlighting their sensitivity to metabolic states; however, their relationship to distinct metabolic complications in obesity remains incompletely understood. To identify the myeloid subpopulations driving the transcriptional responses observed in the myeloid compartment during MASLD progression, we reclustered the myeloid compartment. This resulted in seven distinct subpopulations (Figure 3A and S3A-B), each with a unique marker gene signature (Figure 3B and Table S4). The largest cellular clusters consisted of adipose tissue macrophages (ATMs), lipid-associated macrophages (LAMs) and classical monocytes, while smaller clusters of dendritic cells were also identified (Figure 3A-B). Interestingly, the proportion of LAMs was significantly elevated in patients with steatosis or MASH, whereas other subpopulations did not change with disease stage (Figure 3C and S3C). Accordingly, most LAM-associated neighborhoods displayed increased abundance in patients with steatosis and MASH compared to individuals without MASLD (Figure 3D), also supported by a strong immunofluorescence signal of the LAM-specific marker, TREM2 ^25^, in representative samples from patients with MASLD (Figure 3E and S3D). To further validate the enrichment of LAMs in MASLD, we trained a LASSO regression model ^26^ to predict snRNA-seq-derived LAM proportions based on matched bulk RNA-seq expression of LAM marker genes (Figure S3E). Application of this model to bulk RNA-seq from our larger cohort of female patients revealed higher predicted LAM proportions in SAT from patients with steatosis and MASH than in patients without MASLD (Figure 3F). This is consistent with elevated expression of the LAM markers TREM2 and MMP9 (Figure S3F) and further supports LAM enrichment in SAT of patients with MASLD.

**Figure 3.**
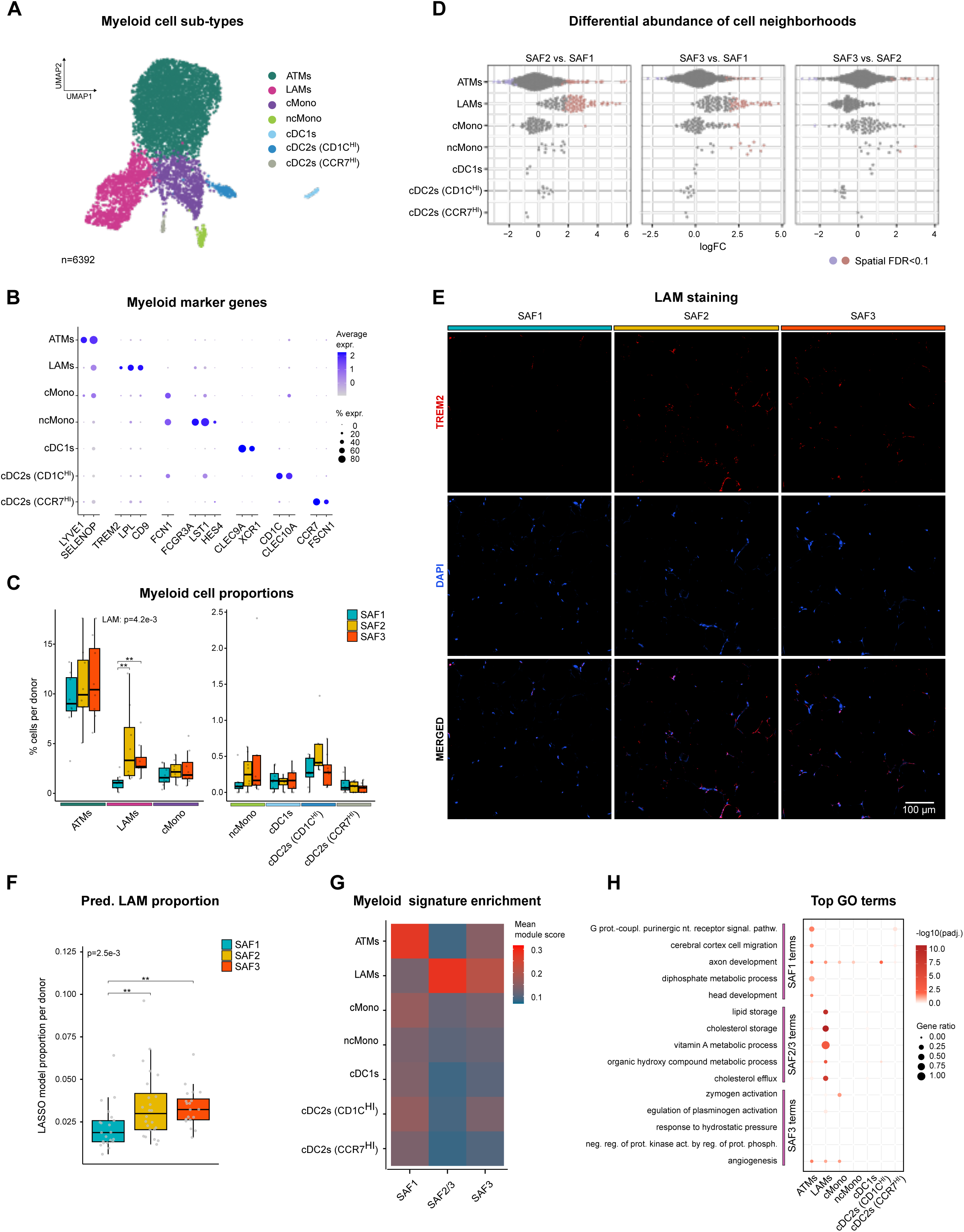
| Lipid-associated macrophages are activated in MASLD. **A**, UMAP representation of myeloid subpopulations from all donors. **B**, Marker genes defining each myeloid subpopulation. **C**, Proportional abundance of myeloid subpopulations in the female SAT snRNA-seq dataset (n=8 per condition). **D**, Distribution of neighborhoods of myeloid subpopulations for indicated SAF score comparisons. Blue, downregulated; Red, upregulated; Grey, unregulated. **E**, Representative immunofluorescent images of TREM2 and DAPI in SAT. **F**, LASSO-based estimation of lipid-associated macrophage (LAM) proportions in the female SAT bulk RNA-seq dataset (S1, n=22; S2, n=25, S3, n=20). **G**, Mean module scores of cluster genes (as defined in Figure 2A) in myeloid subpopulations. **H**, Enrichment of genes linked to top GO biological process pathways (as defined in Figure 2E) in myeloid subpopulations. Boxplot data: center line, median; box limits, upper and lower quartiles; whiskers, 1.5x IQR. A KW test, followed by a post-hoc Wilcoxon rank-sum test with BH correction for multiple testing, was performed to assess significance between SAF diagnosis for **C** and **F** (▪, p<0.1; *, p<0.05; **, p<0.01). ATM, adipose tissue-resident macrophages; cDCs, conventional dendritic cells; cMono, classical monocytes; ncMono, non-classcial monocytes; LAMs, lipid-associated macrophages.

To investigate the contribution of these subpopulations to the overall myeloid transcriptional responses in MASLD, we calculated gene module scores for each of the regulated myeloid gene clusters. The gene module score for the SAF2/3-associated cluster is strongly elevated in LAMs, whereas the SAF1-associated gene cluster is most prominently expressed in ATMs (Figure 3G and S3G). Consistently, genes linked to the SAF2/3-enriched pathways are significantly enriched among LAM marker genes (Figure 3H), reinforcing the role of LAMs in driving most transcriptional reprogramming within the myeloid compartment in MASLD. Interestingly however, ATMs and monocytes also showed increased expression of the lipid-handling gene signature in individuals with steatosis compared to controls without MASLD (Figure S3H). Moreover, we observed a trend toward downregulation of the ATM identity gene program, suggesting both compositional and transcriptional shifts within the myeloid compartment during disease progression (Figure S3H).

In summary, the transition from no-MASLD to MASLD is defined by the expansion of SAT-resident LAMs, the emergence of which drives the observed transcriptional changes in the myeloid immune cell compartment.

### A senescence-associated subpopulation of ASPCs is enriched in patients with MASH

ASPCs have recently been implicated as key contributors to the development of obesity ^27–29^; however, their role in the onset of obesity-related complications, as well as the specific involvement of distinct ASPC subpopulations, remains to be clarified. Subclustering of the ASPC compartment revealed five distinct cellular subclusters, including the previously reported subpopulations *DPP4*^HI^, *CXCL14*^HI^, *PPARG*^HI^, and *EPHA3*^HI^ cells ^16,30^, along with a small but transcriptionally unique cluster, which we designated *PLAU*^HI^ cells (Figure 4A-B and S4A-C). In SAT, *PLAU*^HI^ ASPCs have highly selective expression of several genes encoding signaling molecules, such as *PLAU* itself, *TNFSF14*, *C3*, and *NAMPT* (Figure S4D and Table S5). Interestingly, *PLAU*^HI^ ASPCs have a very low abundance in states of no-MASLD and steatosis but become significantly enriched in the MASH state (Figure 4C and S4C). Accordingly, nearly all *PLAU*^HI^-associated neighborhoods displayed increased abundance in patients with MASH compared to patients with no-MASLD or steastosis (Figure S4E). Interestingly, predicted proportions of *PLAU*^HI^ ASPCs estimated from LASSO regression and PLAU marker gene expression in bulk RNA-seq data vary substantially among patients with MASH in the larger female cohort, indicating marked heterogeneity in *PLAU*^HI^ ASPC abundance in patients with MASH (Figure 4D-E and S4F), which is further addressed below. To evaluate the role of *PLAU*^HI^ ASPCs in driving ASPC transcriptional remodeling, we assessed their contribution to the regulated ASPC gene programs using module scoring. The MASH-linked gene cluster in the ASPCs is strongly associated with *PLAU*^HI^ ASPCs, whereas the SAF1/2-linked gene cluster is more broadly expressed in all other ASPC subpopulations (Figure 4F and S4G), supporting that transcriptional changes in the ASPC compartment in patients with MASH are primarily driven by the emergence of *PLAU*^HI^ ASPCs. Furthermore, several MASH-linked gene programs are enriched in *PLAU*^HI^ ASPCs, including the inflammatory response pathways (Figure S4H). Given that ASPC subpopulations have recently been implicated in cellular senescence in the context of obesity and aging ^23,31^, we investigated whether *PLAU*^HI^ ASPCs might be associated with senescence. Notably, we found that *CDKN1A*, a well-established marker of cellular senescence, was highly enriched in *PLAU*^HI^-associated neighborhoods (Figure 4G). Consistent with a senescent phenotype of *PLAU*^HI^ ASPCs, genes linked to the senescence-associated secretory phenotype (SASP), which is a collection of molecules secreted by senescent cells, were upregulated in these neighborhoods (Figure 4H). Furthermore, the predicted abundance of *PLAU*^HI^ ASPC correlated with *CDKN1A* and SASP expression in bulk RNA-seq data, supporting a link between *PLAU*^HI^ ASPCs and a senescent phenotype in SAT of patients with MASH (Figure 4I-J).

**Figure 4.**
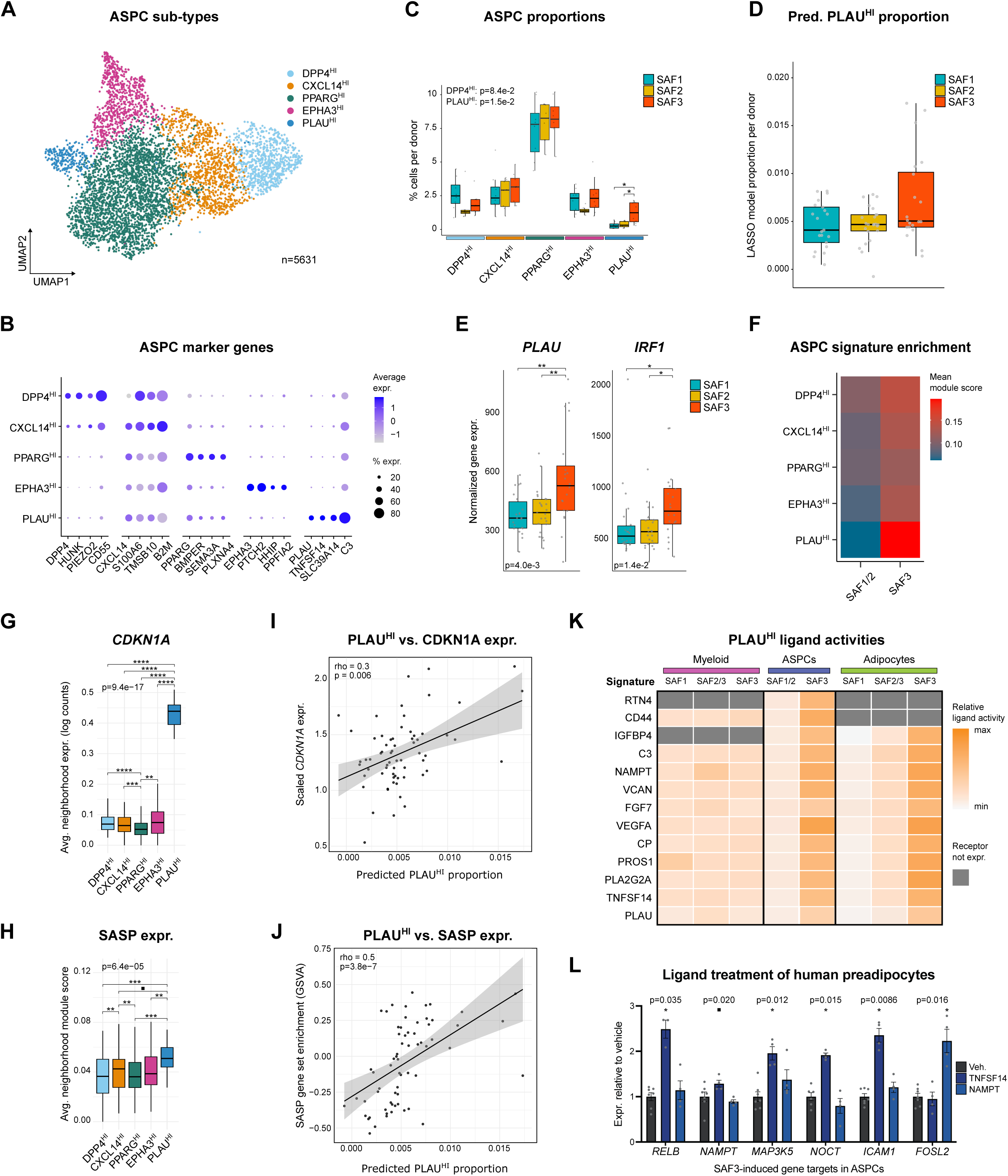
| ASPC remodeling in SAT with increasing MASLD severity. **A**, UMAP representation of ASPC subpopulations from all donors. **B**, Marker genes defining each ASPC subpopulation. **C**, Proportional abundance of ASPC subpopulations in the female SAT snRNA-seq dataset (n=8 per condition). **D**, LASSO-based estimation of *PLAU*^HI^ ASPC proportions in the female SAT bulk RNA-seq dataset (S1, n=22; S2, n=25, S3, n=20). **E**, Normalized expression of selected marker genes of *PLAU*^HI^ ASPCs in the female SAT bulk RNA-seq dataset (S1, n=22; S2, n=25, S3, n=20). **F**, Mean module scores of cluster genes (as defined in Figure 2B) across ASPC subpopulations. **G-H**, Normalized *CDKN1A* (**G)** and SASP marker gene (**H**) expression in indicated ASPC neighborhoods. **I-J**, Correlation between normalized *CDKN1A* (**I**) and SASP marker gene (**J**) expression with LASSO-predicted *PLAU*^HI^ ASPC proportions in 67 female individuals. **K**, Heatmap showing predicted activity of top *PLAU*^HI^ ASPC ligands for regulating cluster genes (as defined in Figure 2A-C) in myeloid immune cells, ASPCs, and adipocytes. **L**, Gene expression analysis in SGBS preadipocytes treated with recombinant TNFSF14 or NAMPT at 20 ng/mL or vehicle control (0.2% BSA) for 24 h. Data represent mean ± SEM (n = 3-7 per condition) and are representative of two independent experiments. Boxplot data: center line, median; box limits, upper and lower quartiles; whiskers, 1.5x IQR. A KW test, followed by a post-hoc Wilcoxon rank-sum test with BH correction for multiple testing, was performed to assess significance between SAF diagnosis for **C-E** or to assess differences between vehicle control and TNFSF14 or NAMPT in **L**. (▪, p<0.1; *, p<0.05; **, p<0.01; ***, p<0.001; ****, p<0.0001). Correlation analyses were performed using Spearman correlation; reported p-values are based on two-sided tests in **I-J**. ASPCs, adipogenic stromal and progenitor cells.

Given that *PLAU*^HI^ ASPCs exhibit a SASP-like state, we investigated whether potential ligands selectively expressed by this population might contribute to MASH-associated transcriptional changes in SAT (Figure S4I). Notably, putative *PLAU*^HI^ ASPC-selective ligands have the highest predicted ligand activity towards MASH-induced gene programs in the ASPCs and adipocytes, suggesting that *PLAU*^HI^ ASPCs may be involved in promoting dysfunction in ASPCs and adipocytes in SAT of patients with MASH (Figure 4K). Consistent with these predictions, ligand stimulation of human preadipocytes with two *PLAU*^HI^ ASPC-associated ligands, TNFSF14 or NAMPT, induced MASH-associated gene targets, supporting a role for these ligands in driving inflammatory and stress-related transcriptional programs in ASPCs (Figure 4L).

In summary, we identified an ASPC subpopulation enriched in female patients with MASH that is associated with a senescent phenotype and exhibits signaling capabilities that may promote SAT dysfunction.

### Loss of a metabolically resilient adipocyte subset from SAT during MASLD progression

Human adipose tissue has recently been shown to harbor transcriptionally distinct adipocyte subpopulations with variable insulin sensitivity ^32^, emphasizing that adipocyte diversity may influence metabolic outcomes. To explore such diversity in relation to MASLD progression and understand how it emerges, we investigated the relationship between adipocyte and ASPC states by mapping trajectories from progenitors to mature adipocytes ^33,34^. This resulted in a strong separation of ASPC and adipocyte nuclei in DC1 and of ASPC subpopulation nuclei in DC2 (Figure 5A and S5A). Interestingly, in the DC1/DC2 plane, two distinct trajectories emerge: one representing adipogenesis and the other representing an alternative specification for the ASPCs (Figure 5A). In accordance with our previous findings ^16,35^, the adipogenic pathway appears to initiate in *DPP4*^HI^ ASPCs, progressing through *CXCL14*^HI^ and *PPARG*^HI^ ASPCs, ultimately culminating in the formation of mature adipocytes (Figure 5A and S5A-C). Neither *PLAU*^HI^ nor *EPHA3*^HI^ ASPCs are a part of the adipogenic trajectory, supporting that these ASPCs may play an alternative, regulatory role in SAT. Further inspection of the adipogenic trajectory and pseudotime analysis revealed a distinct subset of adipocytes (*ADH1B*^HI^) characterized by a unique marker gene signature compared with the remaining (*SFRP4*^HI^) adipocytes (Figure 5B-C and Table S6). *ADH1B*^HI^ adipocytes and their associated neighborhoods are enriched in patients without MASLD relative to those with hepatic steatosis or MASH (Figure 5D and S5D-E). Accordingly, using LASSO regression (Figure S5F), we found that predicted proportions of *ADH1B*^HI^ adipocytes as well as their associated marker genes were significantly higher in SAF1 patients compared to SAF2 and SAF3 patients in the larger cohort of female patients (Figure 5E-F). Moreover, module scores derived from the top marker genes indicated that *ADH1B*^HI^ adipocytes are reduced in obesity in SAT but recover following surgery-induced weight loss ^36^ (Figure 5G).

**Figure 5.**
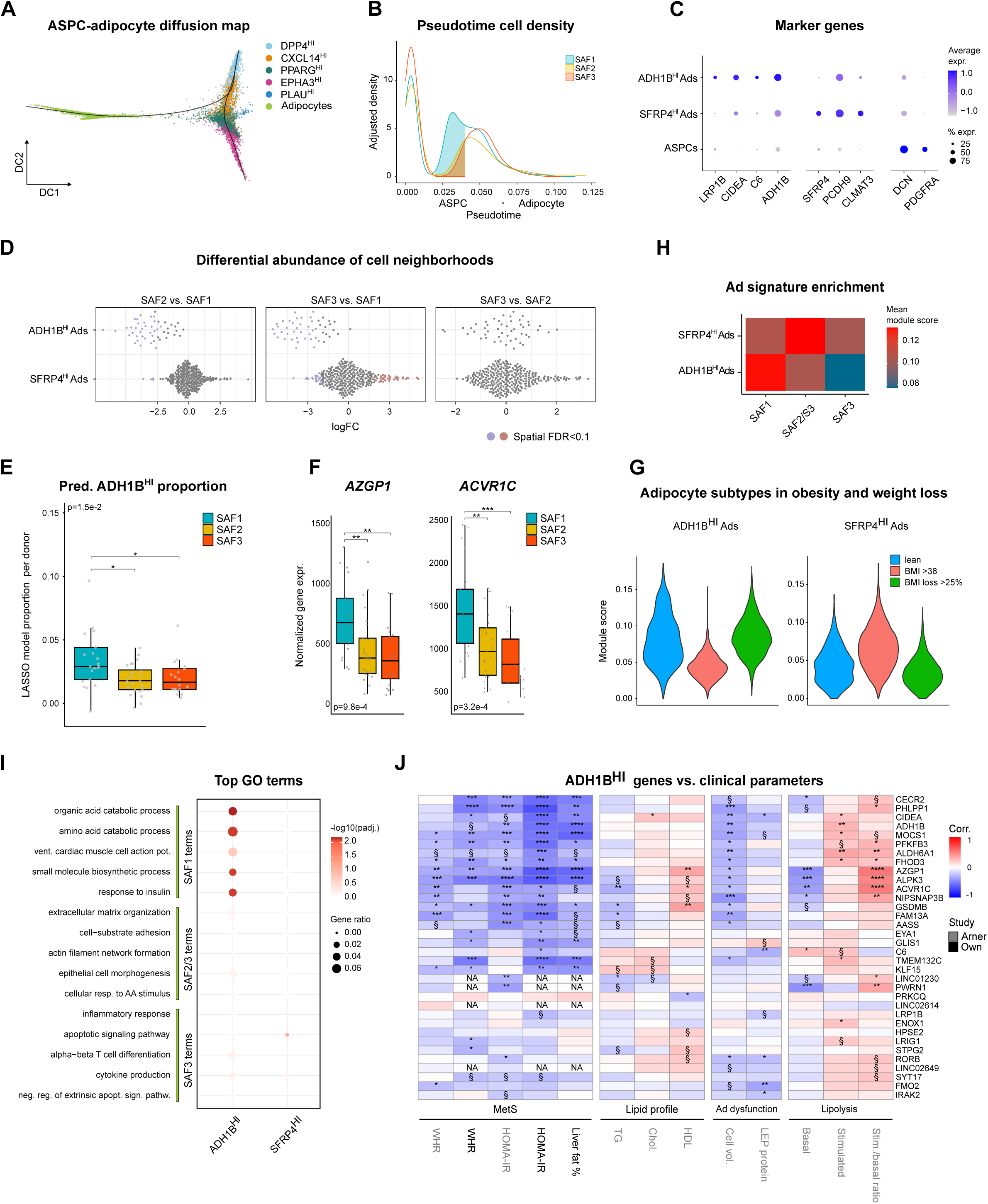
| *ADH1B*^HI^ adipocytes are lost in individuals with MASLD. **A**, Diffusion map (DC1 vs DC2) of ASPC and adipocyte nuclei from all donors, indicating two distinct differentiation trajectories. **B**, Nuclei density across pseudotime, stratified by SAF score. **C**, Marker genes defining adipocyte subpopulations and ASPCs. **D**, Distribution of adipocyte state neighborhoods for indicated SAF score comparisons. Blue, downregulated; Red, upregulated; Grey, unregulated. **E**, LASSO-based estimation of *ADH1B*^HI^ adipocyte proportions in the female SAT bulk RNA-seq dataset (S1, n=22; S2, n=25, S3, n=20). **F**, Normalized expression of selected *ADH1B*^HI^ adipocyte marker genes in the female SAT bulk RNA-seq dataset (S1, n=22; S2, n=25, S3, n=20). **G**, Mean module scores of *ADH1B*^HI^ (left) and *SFRP4*^HI^ (right) adipocyte marker genes in adipocytes from a cohort of lean, obese, and surgery-regressed individuals ^36^. **H**, Mean module scores of cluster genes (as defined in Figure 2C) across adipocyte states. **I**, Enrichment of genes linked to top GO biological process pathways (as defined in Figure 2E) in adipocyte states. **J**, Heatmap showing partial Spearman correlation coefficients between *ADH1B*^HI^ adipocyte marker genes and selected clinical, biochemical, and functional parameters, adjusting for BMI from our cohort (marked in bold) or from Arner et al ^37^. Boxplot data: center line, median; box limits, upper and lower quartiles; whiskers, 1.5x IQR. A KW test, followed by a post-hoc Wilcoxon rank-sum test with BH correction for multiple testing, was performed to assess significance between SAF diagnosis for **E-G**. P-values were calculated using asymptotic methods and corrected for multiple testing using the BH procedure in **J** (▪/§, p<0.1; *, p<0.05; **, p<0.01; ***, p<0.001; ****, p<0.0001).

We next examined how *ADH1B*^HI^ adipocytes relate to the transcriptional reprogramming of the adipocyte compartment in MASLD. We found that the gene module score associated with the SAF1 cluster is elevated in *ADH1B*^HI^ adipocytes (Figure 5H). Consistently, genes involved in SAF1-linked fat metabolism and insulin-sensitizing pathways are significantly enriched among the marker genes of *ADH1B*^HI^ adipocytes (Figure 5I). To further characterize *ADH1B*^HI^ adipocytes, we defined a set of marker genes selectively expressed at the whole-tissue level and correlated these genes with clinical parameters linked to metabolic health status ^37^. Several *ADH1B*^HI^ markers were negatively correlated with indicators of poor metabolic health, including HOMA-IR, WHR, and liver fat (Figure 5J). Furthermore, low expression of a subset of markers such as *AZGP1* and *ACVR1C*, was associated with high basal lipolysis and a low isoprenaline-to-basal lipolysis ratio in adipocytes isolated from SAT ^37^ (Figure 5J), suggesting a potential link between increased basal lipolysis, impaired β-adrenergic responsiveness, and reduced expression of *ADH1B*^HI^ adipocyte marker genes.

Together, these findings suggest that a subset of *ADH1B*^HI^ adipocytes, characterized by improved lipid buffering capacity and linked to enhanced systemic metabolic function, are progressively depleted in obesity, with the greatest loss observed in steatosis and MASH.

### LAMs and *PLAU*^HI^ ASPCs are differently associated with clinical endotypes of MASLD

MASLD and MASH show substantial variation in their prevalence and underlying mechanisms in obesity, with recent findings suggesting at least two distinct clinical forms. One is a liver-specific form, linked to genetic MASLD risk variants and associated with high rates of MASH and progression to chronic liver disease. The other is a cardiometabolic form, occurring independently of genetic risk variants, and characterized by both a high prevalence of MASH and an elevated risk of type 2 diabetes and cardiovascular disease ^4^. To examine how disease-linked subpopulations in SAT link to these different clinical subtypes of MASLD, we clustered patients from an expanded cohort of more than 200 female and male individuals with obesity based on 6 clinical parameters (BMI, age, LDL, ALT, HbA1C, and triglycerides) using a simple clustering algorithm ^4^ (Figure 6A). This resulted in 18.4% (41 patients) and 5.8% (13 patients) of our cohort patients falling into the liver-specific and cardiometabolic MASLD cluster, respectively (Figure S6A). As expected, patients in both these clusters had higher prevalence of MASH and advanced fibrosis compared to the control cluster (Figure 6B), with the liver-specific cluster showing the highest blood levels of the liver damage marker ALT (Table 2). Furthermore, patients in the cardiometabolic cluster had higher serum levels of HbA1C and fasting glucose and were more frequently diagnosed with type 2 diabetes and other metabolic comorbidities compared to both the control and liver-specific cluster (Table 2). We performed bulk RNA-seq of SAT from patients in the expanded cohort and applied our LASSO regression models to estimate the proportions of LAMs, *PLAU*^HI^ ASPCs, and *ADH1B*^HI^ adipocytes across clinical MASLD clusters. Interestingly, predicted LAM abundance and marker gene expression were elevated in both MASH-related endotypes compared to controls (Figure 6C-D). In contrast, *PLAU*^HI^ ASPCs and their markers were specifically enriched in the cardiometabolic MASLD cluster (Figure 6E-F). *ADH1B*^HI^ adipocytes tended to be less prevalent, and their marker genes were downregulated in MASH-related clusters (Figure S6B-C). Together, these findings suggest that LAMs are associated with general adipose tissue dysfunction contributing to hepatic lipid accumulation, whereas *PLAU*^HI^ ASPCs are linked to a more severe adipose tissue and systemic dysfunction.

**Figure 6.**
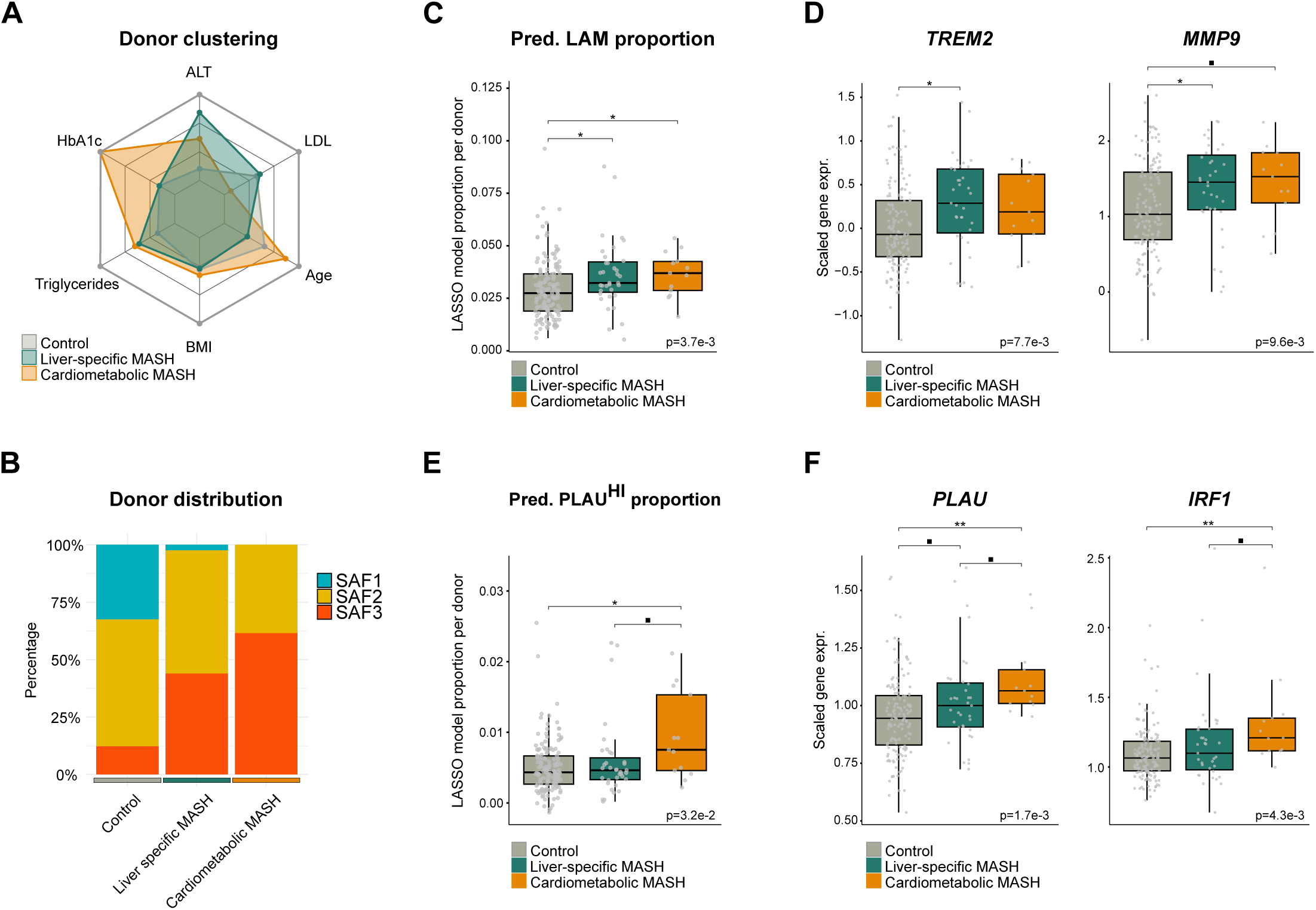
| Adipose tissue subpopulations associate differently with MASH endotypes. **A**, Radar charts showing median values of age, BMI, HbA1c, LDL, triglycerides, and ALT for individuals in the cardiometabolic and liver-specific MASH clusters for the expanded cohort (Control, n=169; LS, n=41, CM, n=13). **B**, Bar charts illustrating the distribution of SAF scores across the indicated clusters. **C**, LASSO-predicted LAM proportions in bulk RNA-seq data in control and MASH endotype clusters (Control, n=146; LS, n=36, CM, n=13). **D**, Normalized expression of selected LAM marker genes from bulk RNA-seq data in control and MASH endotype clusters (Control, n=146; LS, n=36, CM, n=13). **E**, LASSO-predicted *PLAU*^HI^ ASPC proportions in bulk RNA-seq data in control and MASH endotype clusters (Control, n=146; LS, n=36, CM, n=13). **F**, Normalized expression of selected *PLAU*^HI^ ASPC marker genes from bulk RNA-seq data in control and MASH endotype clusters (Control, n=146; LS, n=36, CM, n=13). Boxplot data: center line, median; box limits, upper and lower quartiles; whiskers, 1.5x IQR. A KW test, followed by a post-hoc Wilcoxon rank-sum test with BH correction for multiple testing, was performed to assess significance between SAF diagnosis for **C-F** (▪, p<0.1; *, p<0.05; **, p<0.01). ASPCs, adipogenic stromal and progenitor cells; LAMs, lipid-associated macrophages.

**Table 2.** | Clinical, biochemical, and functional characteristics of extended cohort stratified by clinical subtypes of MASLD. The table indicates mean values (±SD) or % for 223 individuals (158 females and 65 males) stratified into a control cluster and cardiometabolic and liver-specific MASLD endotypes. A KW test, followed by a post-hoc Wilcoxon rank-sum test with BH correction for multiple testing, was performed to assess significance between control group and MASLD endotypes. ALT, alanine aminotransferase; AST, aspartate aminotransferase; BMI, body mass index; CM, cardiometabolic MASLD; Ctrl, control group; HbA1c, glycated hemoglobin A1c; HDL, high-density lipoprotein; LDL, low-density lipoprotein; LS, liver-specific MASLD; MASH, metabolic dysfunction-associated steatohepatitis; TAG, triacylglycerol.

| Variable | Ctrl<br>(n=169) | CM<br>(n=13) | LS<br>(n=41) | P-value<br>(KW) | Ctrl vs LS | Ctrl vs CM<br>(Wilcoxon) | LS vs CM |
| --- | --- | --- | --- | --- | --- | --- | --- |
| Age | 46 (19) | 57 (9) | 37 (13) | <0.001 | 0.0046 | 0.0046 | <0.001 |
| BMI | 42.2 (8) | 43.4 (7.7) | 42 (6.5) | 0.605 | 0.7407 | 0.8594 | 0.7407 |
| HbA1c (mmol/mol) | 37 (7) | 60 (6) | 37 (8) | <0.001 | 0.2862 | <0.001 | <0.001 |
| Fasting blood glucose (mmol/L) | 5.9 (0.9) | 9.7 (2.2) | 6 (1) | <0.001 | 0.4689 | <0.001 | <0.001 |
| Fasting Insulin (mIU/L) | 25.83 (20.25) | 37.83 (10.33) | 41 (36.5) | <0.001 | <0.001 | 0.036 | 0.9378 |
| Total cholesterol (mmol/L) | 4.7 (1.2) | 4.1 (0.4) | 4.8 (1.1) | 0.055 | 0.3937 | 0.049 | 0.049 |
| HDL (mmol/L) | 1.1 (0.4) | 1 (0.2) | 1 (0.2) | 0.0279 | 0.0781 | 0.1236 | 0.724 |
| LDL (mmol/L) | 3.2 (1.1) | 2.4 (0.4) | 3.3 (1) | 0.0018 | 0.4124 | 0.0012 | 0.0012 |
| Blood TAG (mmol/L) | 1.38 (0.8) | 2.16 (0.5) | 2.01 (0.97) | <0.001 | <0.001 | <0.001 | 0.4649 |
| AST (U/L) | 23 (10) | 42 (23.25) | 48 (22) | <0.001 | <0.001 | <0.001 | 0.5466 |
| ALT (U/L) | 27 (17) | 58 (39) | 85 (41) | <0.001 | <0.001 | <0.001 | 0.0442 |
| Antihypertensive drugs (%) | 39.64 | 84.62 | 39.02 | 0.0064 | 1 | 0.0057 | 0.0175 |
| Antidiabetic drugs (%) | 16.57 | 76.92 | 19.51 | <0.001 | 0.7775 | <0.001 | <0.001 |
| Cholesterol-lowering medication (%) | 19.53 | 53.85 | 19.51 | 0.0219 | 1 | 0.0175 | 0.045 |
| Steatosis grade $\geq 1$ (%) | 68.71 | 100.00 | 97.56 | <0.001 | <0.001 | 0.0332 | 1 |
| Ballooning grade $\geq 1$ (%) | 12.88 | 61.54 | 46.34 | <0.001 | <0.001 | <0.001 | 0.6525 |
| Lobular inflammation grade $\geq 1$ (%) | 55.21 | 92.31 | 95.12 | <0.001 | <0.001 | 0.0175 | 1 |
| Fibrosis $\geq 2$ (%) | 19.02 | 61.54 | 48.78 | 0.002 | 0.0332 | 0.0183 | 0.4389 |
| Fibrosis $\geq 3$ (%) | 4.29 | 30.77 | 14.63 | <0.001 | <0.001 | 0.0041 | 1 |
| MASH (%) | 12.27 | 61.54 | 43.90 | <0.001 | <0.001 | <0.001 | 0.4612 |
| Type II Diabetes (%) | 12.43 | 84.62 | 21.95 | <0.001 | 0.1962 | <0.001 | <0.001 |
| Hypertension (%) | 35.71 | 92.31 | 42.50 | <0.001 | 0.6018 | <0.001 | 0.0064 |
| Hyper-cholesterolemia (%) | 18.93 | 61.54 | 20.00 | 0.0037 | 0.9602 | 0.0041 | 0.0204 |

## Discussion

Our findings demonstrate that severe MASLD is associated with specific cellular and molecular signatures of SAT. We observed a reduction in adipocyte fraction alongside hypertrophy, suggesting impaired adipose tissue hyperplasia in MASLD. Concurrently, increased activation of LAMs, along with suppression of key metabolic gene programs in adipocytes and induction of inflammatory and ECM-related gene programs in adipocytes and ASPCs, point to a shift toward a pro-inflammatory and fibrotic tissue environment. These coordinated changes implicate SAT dysfunction as a central contributor to MASLD progression.

Our identification of a transcriptionally distinct subset of *ADH1B*^HI^ adipocytes enriched in individuals with obesity but without MASLD further highlights the importance of adipocyte quality in maintaining systemic metabolic homeostasis. These *ADH1B*^HI^ adipocytes are characterized by elevated expression of gene programs linked to lipid metabolism and insulin sensitization and these are depleted in steatosis and MASH. Notably, some of their marker genes, including *ADH1B*, *AZGP1,* and *ACVR1C*, are inversely associated with clinical indicators of metabolic dysfunction and impaired lipolytic control in adipocytes ^37–39^. This suggests that *ADH1B*^HI^ adipocytes may preserve metabolic flexibility by maintaining appropriate lipid buffering capacity and β-adrenergic responsiveness. In contrast, their depletion during MASLD development may contribute to adipose tissue dysfunction, characterized by unsuppressed basal lipolysis, promoting systemic insulin resistance and MASLD in human obesity. Some *ADH1B*^HI^ adipocyte markers have been functionally implicated in MASLD pathogenesis; AZGP1 has been shown to attenuate MASLD by modulating inflammation and lipolysis ^40,41^, while inhibition of Activin E-ACVR1C signaling in adipose tissue increases lipolysis and hepatic steatosis in mouse models ^42,43^. Together, these findings support a model in which a partial preservation of *ADH1B*^HI^ adipocytes, defined by their transcriptional profile and functional lipid handling, reflects a metabolically resilient adipocyte state that protects against hepatic steatosis and MASH. Their restoration following weight loss further suggests that this adipocyte state is dynamic and potentially therapeutically targetable.

The proportion of LAMs in adipose tissue is elevated in obesity in both mice and humans ^23,25,44–46^ and is gradually reversed toward lean levels following bariatric surgery-induced weight loss ^16,23,47^, highlighting their responsiveness to metabolic state transitions. However, their enrichment in adipose tissue of individuals with varying degrees of metabolic complications has been less well characterized. A recent study reported LAM enrichment exclusively in visceral adipose tissue (VAT) of male individuals, with no corresponding changes in SAT when comparing insulin-sensitive and insulin-resistant individuals with obesity ^48^, suggesting that LAM accumulation may be influenced by disease severity, metabolic phenotype, and sex. Only minor, non-significant shifts in LAM-like cell proportions were observed across MASLD stages in VAT ^13^. In contrast, we found a pronounced enrichment of LAMs in SAT from patients with MASLD compared to individuals without, indicating depot-dependent differences in the adipose tissue response to MASLD. This may reflect a state of SAT dysfunction, where impaired adipocyte lipid storage capacity together with LAM activation drives systemic lipid overflow and ectopic fat accumulation in the liver. Whereas LAM enrichment in SAT is already established at the steatotic stage and remains relatively stable with increasing MASLD severity, TREM2^+^ macrophages in the liver is typically confined to more advanced disease stages with inflammatory and fibrotic changes ^49^. This divergence highlights a tissue-specific timing and threshold for LAM activation, with SAT acting as an early sentinel of metabolic stress. This early adaptation in adipose tissue likely represents a compensatory attempt to manage increased lipid flux and inflammation, which may become maladaptive if sustained.

While senescence in ASPC subpopulations has been implicated in obesity ^23^, our study extends this observation by identifying a transcriptionally distinct *PLAU*^HI^ ASPC subpopulation specifically enriched in patients with MASH. *PLAU*^HI^ ASPCs exhibit a unique combination of SASP-like programs and pro-inflammatory ligand signaling, positioning them as potential senescence-amplifying hubs within SAT. Mechanistically, *PLAU*^HI^ ASPCs express ligands such as TNFSF14 and NAMPT with the potential to activate stress and inflammatory gene programs in ASPCs and adipocytes, indicating a capacity to propagate tissue dysfunction. Persistent senescence in ASPCs may override regenerative cues, contributing to impaired adipogenesis, altered secretory profiles, and chronic inflammation in SAT that may drive metabolic dysfunction ^50–53^. Given that senescence in ASPCs can be reversed by senolytic or SASP-targeting interventions ^54,55^, *PLAU*^HI^ ASPCs may represent a modifiable cellular target for precision therapies in MASLD.

We explored clinical relevance by stratifying patients into liver-specific and cardiometabolic MASLD endotypes ^4^ revealing distinct patterns of adipose remodeling in MASLD. LAMs were elevated in both endotypes, whereas *ADH1B*^HI^ adipocytes tended to be less abundant across MASH-related cluster, suggesting that some degree of adipose tissue dysfunction is required for hepatic steatosis even in the liver-specific cluster. Importantly, *PLAU*^HI^ ASPCs were selectively enriched in the cardiometabolic MASLD cluster, linking this senescent progenitor state to systemic metabolic dysfunction and increased disease severity. This enrichment highlights *PLAU*^HI^ ASPCs as a cellular marker of cardiometabolic risk and a potential therapeutic target in MASLD beyond genetic predisposition.

In summary, our data suggest that adipose tissue remodeling is a key driver of MASLD progression, where macrophage activation, progenitor senescence, and adipocyte dysfunction converge to promote systemic metabolic impairment. These insights may open avenues for precision medicine approaches, particularly in cardiometabolic MASLD characterized by broader systemic involvement.

### Limitations of the study

The snRNA-seq analyses were based on a relatively modest cohort of 24 individuals selected to represent characteristic MASLD archetypes, which may limit detection of subtle or rare cellular states. Nonetheless, key observations from snRNA-seq were validated in bulk RNA-seq from an expanded cohort of ∼200 individuals, supporting their robustness and generalizability. Our analyses were restricted to SAT, so depot-specific features relevant to liver–adipose tissue cross-talk may not be captured. Furthermore, the cross-sectional design constrains causal inference, underscoring the importance of future longitudinal and mechanistic studies to establish the role of identified cell states in MASLD progression.

## Online Methods

### Study participant details

All participants in this study were recruited as part of a clinical cohort previously described in detail ^56,57^, enrolled in a prospective case–control trial of severe obesity (PROMETHEUS). Inclusion criteria were age 18–70 years and BMI ≥30 kg/m². Exclusion criteria included excessive alcohol intake (>12 g/day for women, >24 g/day for men), known or newly diagnosed chronic liver diseases, use of hepatotoxic medications (e.g., glucocorticoids, tamoxifen, amiodarone), short life expectancy, or contraindications to liver biopsy. For the present analysis, we included an initial cohort of 67 female participants and an expanded cohort of 223 individuals (158 females and 65 males).

The PROMETHEUS trial is registered at https://ClinicalTrials.gov (NCT03535142), and was approved by the Regional Committee on Health Research Ethics (S-20170210 and S-20160006G). All participants provided written informed consent prior to enrollment. All procedures complied with relevant ethical regulations and were conducted according to a standardized protocol. This included liver and adipose tissue biopsies, medical history, transient elastography (FibroScan, Echosens, Paris, France), and blood sampling, with data collected on the same day under fasting conditions. Data management was carried out in REDCap via the Odense Patient Data Explorative Network (OPEN.rsyd.dk; OP-551)

### Sampling of human SAT samples

Abdominal SAT biopsies were sampled under sterile conditions by two trained clinicians after a bedside ultrasound-guided marking to determine optimal placement of a liver biopsy. The SAT biopsy site was therefore typically between the seventh and eighth intercostal space at the mid-axillary line. Samples were immediately placed into sterile saline water, divided into smaller pieces and preserved immediately in RNAlater (Sigma-Aldrich, St. Louis, MO) or snap-frozen in liquid nitrogen. Blood samples were collected by an experienced laboratory technician. Biochemical analyses were conducted following standard regional protocols using commercially available kits. All samples were processed by specialized research biochemical technicians and stored at -80 °C.

### Histology and staging of MASLD

The procedure for staging of MASLD was performed essentially as previously described ^57^. In brief, liver biopsies were fixed in 4 % (w/v) neutrally buffered formalin, embedded in paraffin and cut into 4 μm consecutive slices for hematoxylin and eosin (H&E) and Sirius red staining. Biopsies were considered sufficient if ≥10 mm in length with 6 or more portal tracts or regenerative nodules. Liver biopsies were assessed by a single trained pathologist who was blinded to all other data. Scoring followed the NASH Clinical Research Network (NAS-CRN) system for NAFLD, evaluating steatosis (0-3), lobular inflammation (0-3), and ballooning (0-2) ^58^. The NAFLD activity score (NAS) ranged from 0 to 8, calculated as the sum of these components. Fibrosis was graded according to the Kleiner classification: F0 (no fibrosis) to F4 (cirrhosis). Based on the histological assessment, we divided the cohort into three outcome groups: patients with no-MASLD (SAF 1), steatosis (SAF 2), or MASH (SAF 3) using the SAF scoring system ^17^.

### Histology of adipose tissue

#### Adipocyte size

5 µm-sectioned formalin-fixed paraffin-embedded (FFPE) SAT sections were deparaffinized using Tissue-Tek Tissue-Clear (Sakura, cat. no. 1466) for 7.5 min followed by rehydration in graded ethanol solutions and ddH_2_O. Masson’s trichrome staining was performed using the Trichrome Stain Kit (Abcam, cat. no. ab150686) according to manufacturer’s instructions with final mounting in Eukitt® Quick-hardening mounting medium (Sigma-Aldrich, cat. no. 03989). Sections were imaged using a Nikon Ti2 Widefield microscope at 20x magnification and the NIS-Elements AR software (Version 5.30.07). Images of individual regions were stitched together using the NIS-Elements JOBS function to obtain coverage of the whole section area.

#### Immunofluorescent staining of TREM2 and F4/80

5 µm-sectioned FFPE SAT sections were deparaffinized and rehydrated through graded ethanol as described above. Subsequently, sections were subjected to heat-induced antigen retrieval in sodium citrate buffer (pH□6.0) (Sigma-Aldrich, cat. no. C9999) for 30□min. Sections were permeabilized with 0.3% Triton X-100 (Sigma-Aldrich, cat. no. X100) and blocked in PBS containing 1% donkey serum (Jackson ImmunoResearch, cat. no. 017-000-121) and 1% BSA (Sigma-Aldrich, cat. no. A9647) for 30□min at room temperature. Primary antibodies against TREM2 (R&D Systems, cat. no. MAB17291, rat, 1:50) and F4/80 (Proteintech, cat no. 28463-1-AP, rabbit, 1:250) were applied overnight at 4□°C, followed by incubation with anti-rat IgG Alexa Fluor 647 secondary antibody (Invitrogen, cat. no. A78947, 1:500), anti-rabbit IgG Alexa Fluor 568 secondary antibody (Invitrogen, cat. no. A10042, 1:500) and DAPI (Sigma Aldrich, cat. no. D9542, 1:500) for 1□h at room temperature in the dark. Slides were mounted with ProLong Diamond Antifade Mountant (Invitrogen, cat. no. P36965) and imaged using a Nikon A1 confocal microscope (20× objective) and NIS-Elements AR software. Negative controls included blank and secondary-only staining.

### RNA extraction

SAT was homogenized in TRI Reagent® (Sigma-Aldrich) using the FastPrep-24™ instrument (MP Biomedicals) followed by RNA extraction using EconoSpin columns (Epoch). NEBNext Ultra RNA Library Prep Kit for Illumina (New England Biolabs, San Diego, CA) was used to construct libraries according to the manufacturer’s protocol. RNA libraries were paired-end-sequenced using the NovaSeq™ 6000 platform (Illumina, San Diego, CA).

### Nuclei Isolation

Nuclei from SAT were processed in two experimental pools for single nuclei sequencing. The first pool for snRNA-seq, nuclei were isolated by first coarsely mincing each tissue piece on dry ice, followed by homogenization in 2 mL of Nuclei Extraction Buffer (Miltenyi Biotec, 130-128-024) supplemented with 0.5 U/mL RNase inhibitor (New England Biolabs, M0314) using the 4C_nuclei_1 program on the gentleMACS^TM^ Octo Dissociator (Miltenyi Biotec, 130-096-427) with gentleMACS^TM^ C Tubes (Miltenyi Biotec, 130-093-237). The resulting nuclei suspension was filtered through a 70 µm MACS® SmartStrainer (Miltenyi Biotec, 130-110-916) and centrifuged at 500×g at 4 °C for 10 min. The nuclei pellet was resuspended in 80 µL nuclei resuspension buffer (1 mg/mL BSA (Sigma-Aldrich, B6917), 0.04 U/μL RNase inhibitor (New England Biolabs, M0314), 1x PBS (Gibco, 70013-016) in DEPC-treated water) and filtered through a pre-wetted 40 µm Flowmi® Cell Strainer (Merck, BAH136800070). The original tube was washed with an additional 60 µL of nuclei resuspension buffer, which was also filtered through the same strainer, bringing the total volume to 140 µL. Equal numbers of nuclei from each sample were pooled, centrifuged at 500×g at 4 °C for 10 min, and resuspended in 125 µL of nuclei resuspension buffer. Three replicates of 40,000 pooled nuclei were collected for sequencing.

For the second experimental pool, nuclei were isolated following the same initial steps as described above, up to and including the initial centrifugation. The nuclei pellet was resuspended in 100 µL of nuclei resuspension buffer (1 mg/mL BSA (Sigma-Aldrich, B6917), 0.04 U/μL RNase inhibitor (New England Biolabs, M0314), 1x PBS (Gibco, 70013-016) in DEPC-treated water) and centrifuged at 500×g at 4 °C for 10 min. The nuclei pellet was then resuspended in 100 µL of permeabilization buffer (10 mM Tris-HCl (pH 7.5) (Sigma-Aldrich, T2194), 3 mM MgCl_2_ (Sigma-Aldrich, M1028), 10 mM NaCl (Sigma-Aldrich, 59222C), 0.01 % Tween-20 (Sigma-Aldrich, P1379), 0.01 % IGEPAL CA-630 (Sigma-Aldrich, I8896), 1 mg/mL BSA (Sigma-Aldrich, B6917), 0.001 % Digitonin (Invitrogen, BN2006), 1 mM DTT (New England Biolabs, B1034A), 1 U/µL RNase inhibitor (New England Biolabs, M0314), in DEPC-treated water), incubated on ice for 2 min, and subsequently diluted with 1 mL wash buffer (10 mM Tris-HCl (pH 7.5) (Sigma-Aldrich, T2194), 3 mM MgCl_2_ (Sigma-Aldrich, M1028), 10 mM NaCl (Sigma-Aldrich, 59222C), 0.01 % Tween-20 (Sigma-Aldrich, P1379), 1 mg/mL BSA (Sigma-Aldrich, B6917), 1 mM DTT (New England Biolabs, B1034A), 1 U/µL RNase inhibitor (New England Biolabs, M0314) in DEPC-treated water). The solution was centrifuged at 500×g at 4 °C for 10 min, and the nuclei pellet was resuspended in 75 µL of diluted nuclei buffer (1x Nuclei Buffer (10x Genomics, 2000153/2000207), 1 mM DTT (New England Biolabs, B1034A), 1 U/µL RNase inhibitor (New England Biolabs, M0314), in DEPC-treated water) and filtered through a pre-wetted 40 µm Flowmi® Cell Strainer (Merck, BAH136800070). Equal numbers of nuclei from each sample were pooled, centrifuged at 500×g at 4 °C for 10 min, and resuspended in 12 µL of diluted nuclei buffer. Four aliquots of pooled nuclei containing 12,000, 16k,000, 20,000, and 30,000 nuclei, respectively, were collected for sequencing.

### Single-nucleus library preparation and sequencing

Immediately following nuclei isolation, the nuclei were loaded onto the 10x Genomics Chromium controller (10x Genomics, PN110203). For the first experimental pool, libraries were prepared following the manufacturer’s instructions using the 10x Genomics Chromium Next GEM Single Cell 3’ Reagents Kits v3.1 (Dual Index) (User Guide CG000315 Rev E), using Next GEM Single Cell 3’ Gel Beads v3.1 (10x Genomics, PN-2000164), Chromium Next GEM Chip G (10x Genomics, PN-2000177), Chromium Next GEM Single Cell 3’ GEM Kit v3.1 (10x Genomics, PN-1000123), Library Construction Kit (10x Genomics, PN-1000190), and Dual Index Kit TT Set A (10x Genomics, PN-1000215). For the second experimental pool, libraries were prepared following the manufacturer’s instructions using the RNA part of the 10x Genomics Chromium Next GEM Single Cell Multiome ATAC + Gene Expression Kits (Dual Index) (User Guide CG000338 Rev E), using Next GEM Single Cell Multiome Gel Beads A (10x Genomics, PN-2000261), Chromium Next GEM Chip J (10x Genomics, PN-2000264), Chromium Next GEM Single Cell Multiome GEM Kit A (10x Genomics, PN-1000232), Chromium Next GEM Single Cell Multiome Amp Kit A (10x Genomics, PN-1000233), Library Construction Kit (10x Genomics, PN-1000190), Dual Index Kit TT Set A (10x Genomics, PN-1000215)). Sequencing was performed using the NovaSeq™ 6000 platform (Illumina, San Diego, CA).

#### Cell culture and ligand treatment

SGBS preadipocytes were cultured in DMEM/F12 (Gibco, 11320033) supplemented with 10% fetal bovine serum (FBS), 1% penicillin–streptomycin, 2 mM L-glutamine, 33 μM biotin, 17 μM pantothenate, 90 μg/mL heparin, and 1 ng/mL fibroblast growth factor 1 (FGF-1) at 37 °C in 5% CO□. Cells were passaged at 70-80% confluence and seeded at equal density for experiments. After reaching confluence, cultures were maintained for 2 additional days, followed by 24 h of serum starvation. Cells were then treated with recombinant NAMPT (ThermoFisher, 130-09) or TNFSF14 (MedChemExpress, HY-P72522) at 20 ng/mL for 24 h. Vehicle controls received 0.2% BSA in ddH□O.

#### Quantification and statistical analysis

No statistical methods were used to pre-determine sample sizes, but our sample sizes are similar to those reported in a previous snRNA-seq publication ^36^. Statistical analyses were performed in R (v4.4.2) using functions from the *rstatix* and *purrr* packages. Overall differences across SAF conditions were assessed using the Kruskal-Wallis test. If significant, pairwise comparisons between individual SAF groups were performed using Wilcoxon rank-sum tests, with p-values adjusted for multiple testing using the Benjamini-Hochberg (BH) correction. All post hoc tests were two-sided. Correlations were assessed using Pearson’s or Spearman’s correlation coefficient, as indicated. Associations between gene expression of *ADH1B*^HI^ adipocytes and clinical variables (e.g., WHR, HOMA-IR, liver fat percentage) were evaluated using partial Spearman correlation tests, adjusting for BMI. Significance was determined using asymptotic p-values from the partial correlation test, with multiple testing correction applied using BH adjustment. Further details on specific statistical tests are provided in the figure legends. Most plots, including tile plots, boxplots, and dimensionality reductions, were generated directly or indirectly using ggplot2. In boxplot visualizations, each patient is represented as an independent replicate.

### Adipocyte size quantification

Images were white-balanced using Fiji software (ImageJ version 1.54p) and a publicly available ImageJ macro ^59^, followed by conversion into binary masks, denoising and structure closing. Adipocyte area size was finally quantified using particle analysis in Fiji, with area thresholds of 500-56,300 µm^2^ ^60,61^ and circularity thresholds of 0.5-1.0. Only patients with ≥500 quantified adipocytes were included in subsequent analyses.

### Analysis of bulk RNA-seq data

The quality of raw sequencing reads was assessed using *FastQC* (v.0.11.9) ^62^ and *MultiQC* (v1.10.1) ^63^. Raw bulk RNA-seq data were aligned to the human genome assembly (GRCh38, Ensembl release 101) using *STAR* (version 2.7.11a) ^64^. Aligned reads were sorted and indexed using *Samtools* (version 1.17) ^65^ and low-quality reads were filtered out (MAPQ < 30, SAM flag = 780). Exon reads were quantified using *FeatureCounts* (version 2.0.6) ^66^. Initial batch correction was performed using the ComBat-Seq function from the *sva* package (v3.56.0) ^67^ to adjust for differences between sequencing rounds while preserving biological variation across SAF-diagnosed groups.

### Differential gene expression analyses

Differential gene expression analysis, for both all females and the subset of archetype females, was performed using three independent methods: *DESeq2* (v1.48.1) ^68^, *edgeR* (v4.6.3) ^69^, and *dearseq* (v1.20.0) ^70^. In *DESeq2*, batch-corrected counts were modeled to test for differences between SAF groups, adjusting for BMI and age, using the Wald test with independent filtering and multiple testing correction. *edgeR* ^69^ was applied with TMM normalization, dispersion estimation, and quasi-likelihood F-tests, adjusting for BMI and age. *dearseq* ^70^ was applied using the asymptotic test with multiple comparison correction. Only genes identified as differentially expressed (padj < 0.1) by all three methods were retained for downstream analyses, including k-means clustering and functional enrichment analyses using *clusterProfiler* ^71^ and GO biological pathways (v4.16.0).

### Linear discriminant analyses and archetype selection

Linear discriminant analysis (LDA) was performed using *MASS* (v7.3-65) on TMM-normalized bulk RNA-seq data. The 500 most variable genes were selected based on variance across samples, then scaled and centered prior to analysis. Archetype analysis was applied to the resulting LDA projections (LD1 and LD2) using the *archetypes* packages (v.2.2-0.2) ^72^. The optimal number of archetypes (k = 3) was selected by scree plot inspection. Samples with the highest archetypal coefficients were considered representative of each SAF diagnostic group and selected for single-nucleus RNA sequencing.

### Analyses of single-nucleus RNA-seq data

Raw snRNA-seq data were aligned using *Cell Ranger* v. 7.1.0 ^73^ with the GRCh38-2020-A reference. Samples were demultiplexed using bulk RNA-seq data from the included donors. Bulk and single-nucleus RNA-seq data were aligned using *STAR*^64^. Valid single-nucleus barcodes were identified with *valiDrops* ^74^, and donor SNPs were called using *Cellsnp-lite* ^75^. Demultiplexing was performed using *vireo* ^76^. Filtering and QC were performed using *CRMetrics*(v.0.3.2) ^77^ with a depth threshold of 1,000. Mitochondrial and ribosomal gene counts were removed from the count matrices. Cells with high ambient RNA content were identified and removed using *CellBender* within *CRMetrics* ^77^. Doublets were identified using *DoubletDetection*.

Data normalization and scaling (*SCTransform* for global analysis and *ScaleData* for subclusters), PCA, UMAP embedding, shared nearest neighbor construction, and clustering were performed using *Seurat* (v.5.2.1) ^78^. Batch correction between experimental pools (but not across donors) was performed using *Harmony* (v. 1.2.3) ^79^. Cell types and subpopulations were annotated based on established adipose tissue cell type markers ^18^. A *PLAU*^HI^ ASPC population did not match any previously described subtype and was therefore annotated as a novel subpopulation.

Deconvolution to estimate cell type proportions from bulk RNA-seq data was performed using *BayesPrism/TED* (v.1.4) ^80^. DEGs were identified using *Cacoa* (v.0.4.0) ^81^. Heatmaps of per-donor pseudobulk DEGs were generated using *Seurat*, clustered with the built-in *kmeans* function, and visualized with *Pheatmap* (v.1.0.12). All module scores were calculated using *UCell* (v. 2.8.0) ^82^.

Enrichment of ATM, LAM, and monocyte markers across SAF scores was assessed by selecting subcluster marker genes (adjusted P < 1e-50, log2 fold change > 1, pct. expressed > 0.05, pct. expressed in control < 0.5). Module scores derived from these marker sets were assigned to cells, aggregated per donor, SAF score, and subcluster (pseudobulk), and visualized as pseudo bulked module scores.

Gene ontology analysis and subsequent simplification of GO biological process terms were performed using *ClusterProfiler* (v.4.12.6) ^71^. P-values were adjusted using Benjamin Hochberg (BH) correction after simplification. To assess enrichment of SAF signature-associated, cell type-stratified GO terms in subpopulation-specific marker genes, annotated genes for each GO term were first extracted. These gene sets were then tested for overrepresentation among the top 200 most significant marker genes of each cellular subcluster or substate, as identified with *FindMarkers* from *Seurat* ^78^, using Fisher’s exact test.

The Milo R package (version 2.0.0) ^19^ was used to identify cell neighborhoods and assess differential cell type and -state abundance based on KNN graphs. The KNN graph was build using 40 nearest-neighbors (k=40) and 20 dimensions (d=20). Cell neighborhoods were defined on the KNN graph using random sampling of 20% of all cells along with the graph sampling scheme. Cells in neighborhoods were counted per donor followed by specification of contrast of interest (SAF1, SAF2, SAF3) for testing of differential cell abundance using TMM normalization and the graph-overlap spatial FDR weighting scheme. Neighborhoods consisting of ≥70% of the same cell type or -state were included in the analysis and were considered significant if their spatial FDR□<□0.1.

To estimate the proportion of selected cell subtypes or substates in bulk RNA-seq data we used a LASSO regression framework implemented in *glmnet* (v. 4.1.8) ^26^. As input, we used marker genes identified by *FindMarkers* from *Seurat* and adjusted cutoffs based on relevant parameters (p_val_adj, avg_log2FC, pct.2, pct.1) iteratively selecting for optimized performance. Performance was evaluated using leave-one-out cross-validation (LOOCV), and model accuracy was quantified using R and mean absolute error. The final model was trained on all samples using the optimal regularization parameter (lambda) obtained from *cv.glmnet*. Selected genes were extracted from the fitted LASSO model and applied to scaled bulk RNA-seq expression data for full-cohort prediction.

The regulatory potential of PLAU+ ASPC secreted ligands towards myeloid cells, ASPCs, and adipocytes in different MASLD stages was predicted using the *NicheNet* package (v. 2.1.5) ^83,84^ in R (v.4.4.1), along with its associated ligand-receptor network database and ligand-target prior model downloaded from https://zenodo.org/record/7074291/files/lr_network_human_21122021.rds and https://zenodo.org/record/7074291/files/ligand_target_matrix_nsga2r_final.rds, respectively.

Potential ligands were found among PLAU+ ASPC marker genes that had been identified in both the ASPC sub-analysis (Padj.<0.1 and log2FC>0.5) as well as in the global SAT cell atlas analysis (Padj.<0.05). Ligands whose receptors were expressed by ≥5% of the receiving cell type population (i.e. myeloid cells, ASPCs or adipocytes) were included in the analysis.

The gene sets of interest, likely to be impacted by ligand-receptor interactions in each cell type and disease stage were defined as myeloid-, ASPC- and adipocyte-specific disease gene signatures previously identified through cell type-specific DEG analyses across SAF scores and gene clustering (Figure 2A-C). Finally, NicheNet was used to rank ligands based on the similarity between 1) a ligand’s target gene pattern in the prior model and 2) the enrichment of the gene set of interest relative to a background gene set, which was defined as all genes expressed in ≥5% of the receiving cell type.

Pseudo time trajectories were estimated using slingshot based on diffusion map embeddings generated with a combination of *destiny* ^34^, *Seurat* ^78^ and *harmony* ^79^. *ADH1B*^HI^ and *SFRP4*^HI^ adipocytes were defined as an interval in pseudo time, which was selected after iterating over several candidate intervals and evaluating which interval yielded the clearest and most distinct marker gene expression profile.

### MASLD endotypes

A total of 297 patients in our cohort of patients with severe obesity were clustered according to the original algorithm ^4^ using the online tool available at https://ulr-metrics.univ-lille.fr/masldclusters/. Of these, 223 patients were assigned to clusters; 169 patients were assigned to the control cluster, 13 to the cardiometabolic MASLD cluster and 41 to the liver-specific MASLD cluster.

### Public Data

Human abdominal SAT single-nucleus transcriptomic data from the NEFA cohort ^36^ were kindly provided by Laura Hinte and Ferdinand Von Meyenn. Spearman correlation analyses between *ADH1B*^HI^ adipocyte markers and clinical, biochemical, and functional parameters were performed via the Adipose Tissue Knowledge Portal ^85^ using a cohort of female individuals in whom subcutaneous adipocytes had been isolated and assessed for basal and stimulated lipolysis ^37^.

## Data availability

snRNA-seq objects will be made publicly available at the Single Cell Portal at the time of publication under accession number: SCP3208. The raw bulk RNA-seq and snRNA-seq data sets generated in this study cannot be deposited in a public repository due to national data protection laws and restrictions imposed by the ethics committee to ensure participant privacy. However, researchers can apply for access through an individual data processing agreement with the principal clinical investigator (M.M.E.L.) at the University Hospital of Southern Denmark, Esbjerg.

## Code availability

All analyses were conducted using publicly available software packages. The custom code used for RNA-seq data analyses, as well as the datasets generated and/or processed in this study, will be made available on Github upon publication.

## Acknowledgments

We thank Dr. Manbir Sandhu and Dr. Madan Badu (St. Jude Children’s Research Hospital) for expert advice on archetype analysis. We are grateful to Lea Grønkjær and Birgitte Jacobsen (University Hospital of Southern Denmark, Esbjerg) for technical assistance with histological samples. We also thank Dr. Ronni Nielsen at the Functional Genomics and Metabolism Unit (FGM, University of Southern Denmark) for library preparation and sequencing support. SGBS cells were generously provided by Dr. Martin Wabitsch at the University of Ulm, Germany. Confocal fluorescence microscopy was performed at the Danish Molecular Biomedical Imaging Center (DaMBIC, University of Southern Denmark), supported by the Novo Nordisk Foundation (NNF) (NNF18SA0032928). Light-sheet fluorescence microscopy was performed at the Danish Spatial Imaging Consortium (DanSIC, University of Southern Denmark), supported by the NNF. This work was supported by grants from the Danish National Research Foundation to the Center for Functional Genomics and Tissue Plasticity (ATLAS) (Project grant: 141), Sygeforsikringen “Danmark” (grant no. 2021-0297), and the Lundbeck Foundation (R413-2022-471). All computations for this project were performed on the UCloud interactive HPC system, managed by the eScience Center at the University of Southern Denmark.

## Author information

### Authors and Affiliations

**Center for Functional Genomics and Tissue Plasticity (ATLAS), Department of Biochemistry and Molecular Biology, University of Southern Denmark (SDU), Odense, Denmark**

Johanne Drewsen Madsen, Trine Vestergaard Dam, Jyotsna Nambiar, Chunyu Liang, Aino Rakel Peltonen, Louise Søgaard Møller Hansen, Rasmus Rydbirk, Lukas Mc Kenzie Oussoren, Kamilla Schlippe Jakobsen, Viktor Manuel Grunddal Larsen, Ellen Gammelmark Klinggaard, Luna Isabella Wassini, Freja Hald Højgaard, Daniel Hansen, Babukrishna Maniyadath, Kim Ravnskjær, Susanne Mandrup, Anne Loft, Søren Fisker Schmidt.

**The Novo Nordisk Foundation Center for Genomic Mechanisms of Disease, Broad Institute of MIT and Harvard, Cambridge, MA, 02142, USA**

Trine Vestergaard Dam, Susanne Mandrup, Anne Loft, Søren Fisker Schmidt

**Department of Gastroenterology and Hepatology, University Hospital of Southern Denmark, Esbjerg,** Denmark.

Elise Jonasson, Charlotte Wilhelmina Wernberg, Mette Enok Munk Lauridsen

**Department of Pathology, Odense University Hospital, Odense, Denmark**

Tina Di Caterino

**Center for Liver Research (FLASH), Department of Gastroenterology and Hepatology, Odense University Hospital, Odense, Denmark.**

Aleksander Krag

**Institute of Clinical Research, University of Southern Denmark, Odense, Denmark**

Aleksander Krag

## Contributions

Conceptualization and study design were carried out by J.D.M., T.V.D., A.L. and S.F.S. Data acquisition was performed by J.D.M., T.V.D., J.N., C.L., A.R., C.W.W., L.S.M.H., R.R., L.M.O., K.S.J., V.M.G.L., E.G.K., L.I.W., F.H.H., D.H., B.M., E.J., T.D.C., A.K., K.R., S.M, and M.E.M.L. Data analysis and interpretation were conducted by J.D.M., T.V.D., A.L., and S.F.S. Supervision was carried out by A.L., and S.F.S.

---

**Figure S1.**
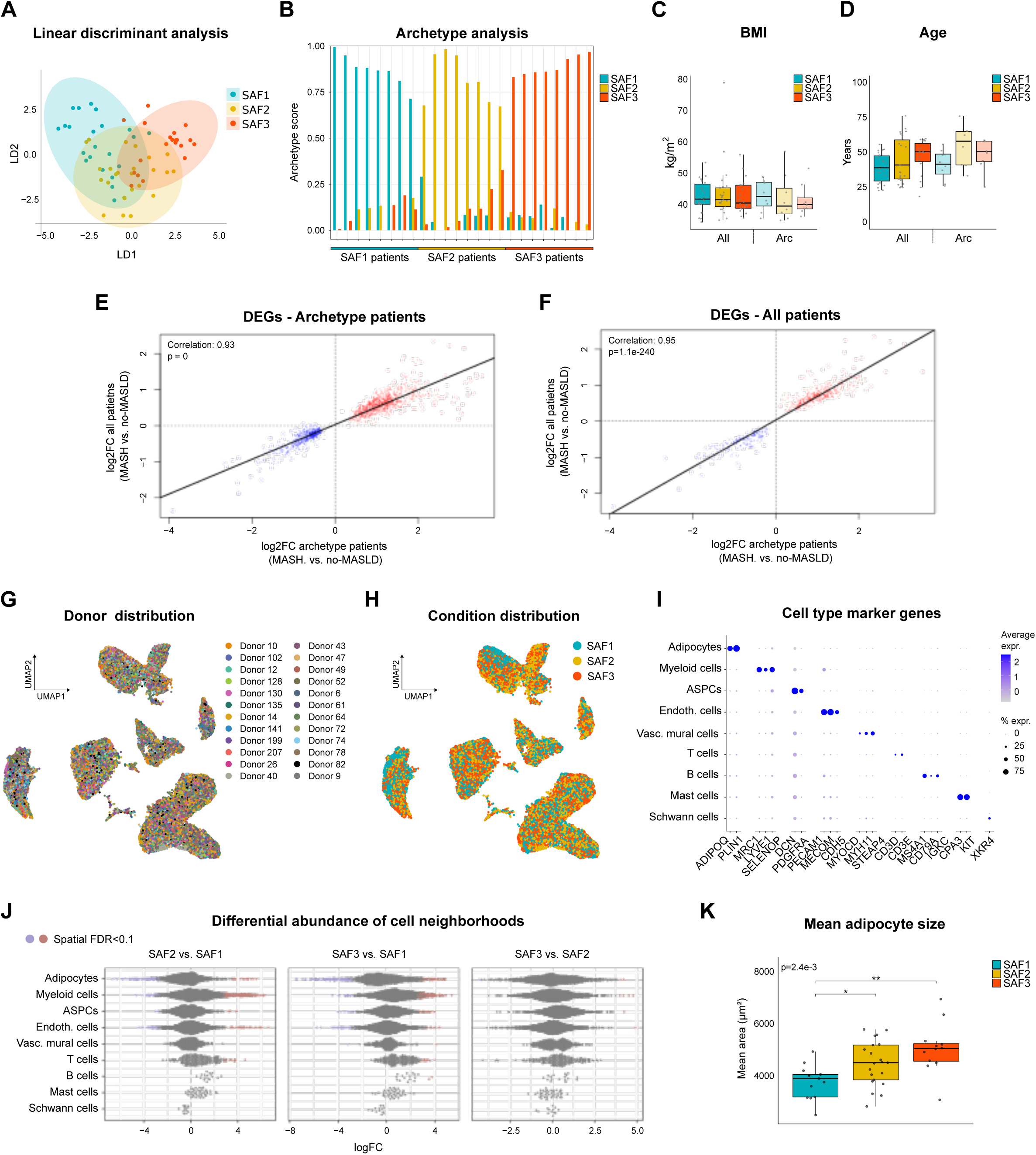
| Transcriptomic and histological profiling of females with obesity stratified by MASLD severity. **A**, Linear discriminant analysis of top 500 most variable genes in the bulk RNA-seq dataset of 67 female individuals. **B**, Archetype scores for each female donor selected for snRNA-seq. **C–D**, Body mass index (**C**) and age (**D**) distribution (all: S1, n=22; S2, n=25, S3, n=20; archetype, n=8 per condition). **E–F**, Correlation between log2 fold change (log2FC) in SAT gene expression (MASH vs no-MASLD) for archetype donors versus all 67 donors, using genes differentially expressed in archetype donors (**E**) and in the full cohort (**F**). **G–H**, UMAP visualization of SAT nuclei from all donors, colored by individual donor (**G**) or SAF score (**H**). **I**, Marker genes defining each cell population in the human SAT snRNA-seq dataset. **J**, Distribution of SAT neighborhoods for the indicated SAF score comparisons. Blue, downregulated; Red, upregulated; Grey, unregulated. **K**, Quantification of mean adipocyte area in SAT (S1, n=13; S2, n=21, S3, n=12). Boxplot data: center line, median; box limits, upper and lower quartiles; whiskers, 1.5x IQR. A KW test, followed by a post-hoc Wilcoxon rank-sum test with BH correction for multiple testing, was performed to assess significance between SAF diagnosis for **C-D** and **K** (▪, p<0.1; *, p<0.05; **, p<0.01). Correlation analyses were performed using Pearson correlation; reported p-values are based on two-sided tests in **E-F**. ASPCs, adipogenic stromal and progenitor cells.

**Figure S2.**
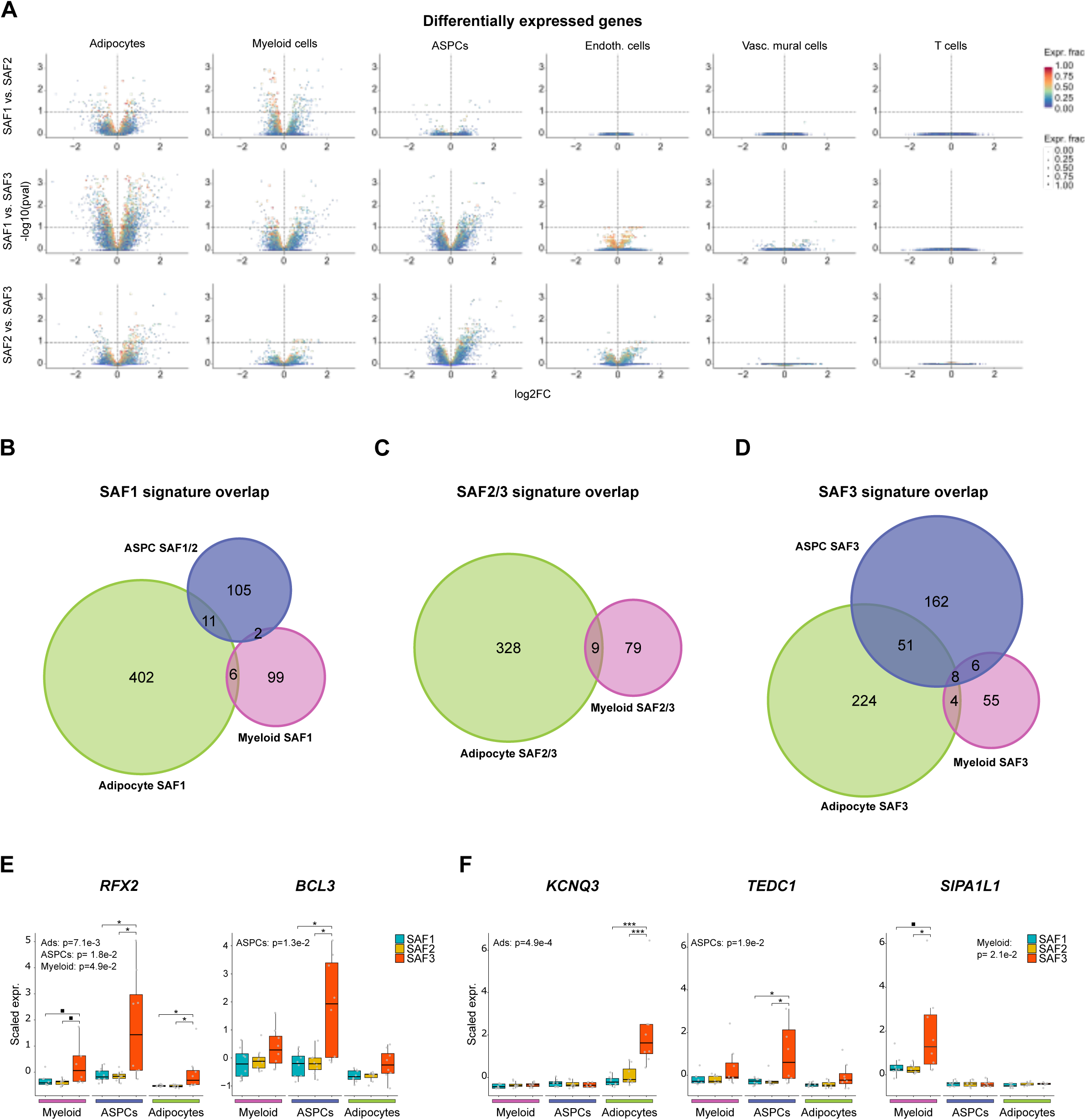
| Cell type-specific transcriptomic signatures in SAT associated with MASLD severity. **A**, Volcano plot showing log₂ fold change in gene expression and distribution of adjusted p-values for indicated SAF score comparisons in major SAT cell types. Genes above the dashed line (padj < 0.1) are considered differentially expressed. **B–D**, Venn diagrams illustrating overlap of cluster genes (as defined in Figure 1A) for SAF1-associated clusters (**B**), SAF2/3-associated clusters (**C**), and SAF3-associated clusters (**D**) for myeloid immune cells, ASPCs, and adipocytes. **E–F**, Normalized pseudobulk expression of shared (**E**) and cell type-specific (**F**) SAF3-associated cluster genes (n=8 per condition). Boxplot data: center line, median; box limits, upper and lower quartiles; whiskers, 1.5x IQR. A KW test, followed by a post-hoc Wilcoxon rank-sum test with BH correction for multiple testing, was performed to assess significance between SAF diagnosis for **E-F** (▪, p<0.1; *, p<0.05; ***, p<0.001). ASPCs, adipogenic stromal and progenitor cells.

**Figure S3.**
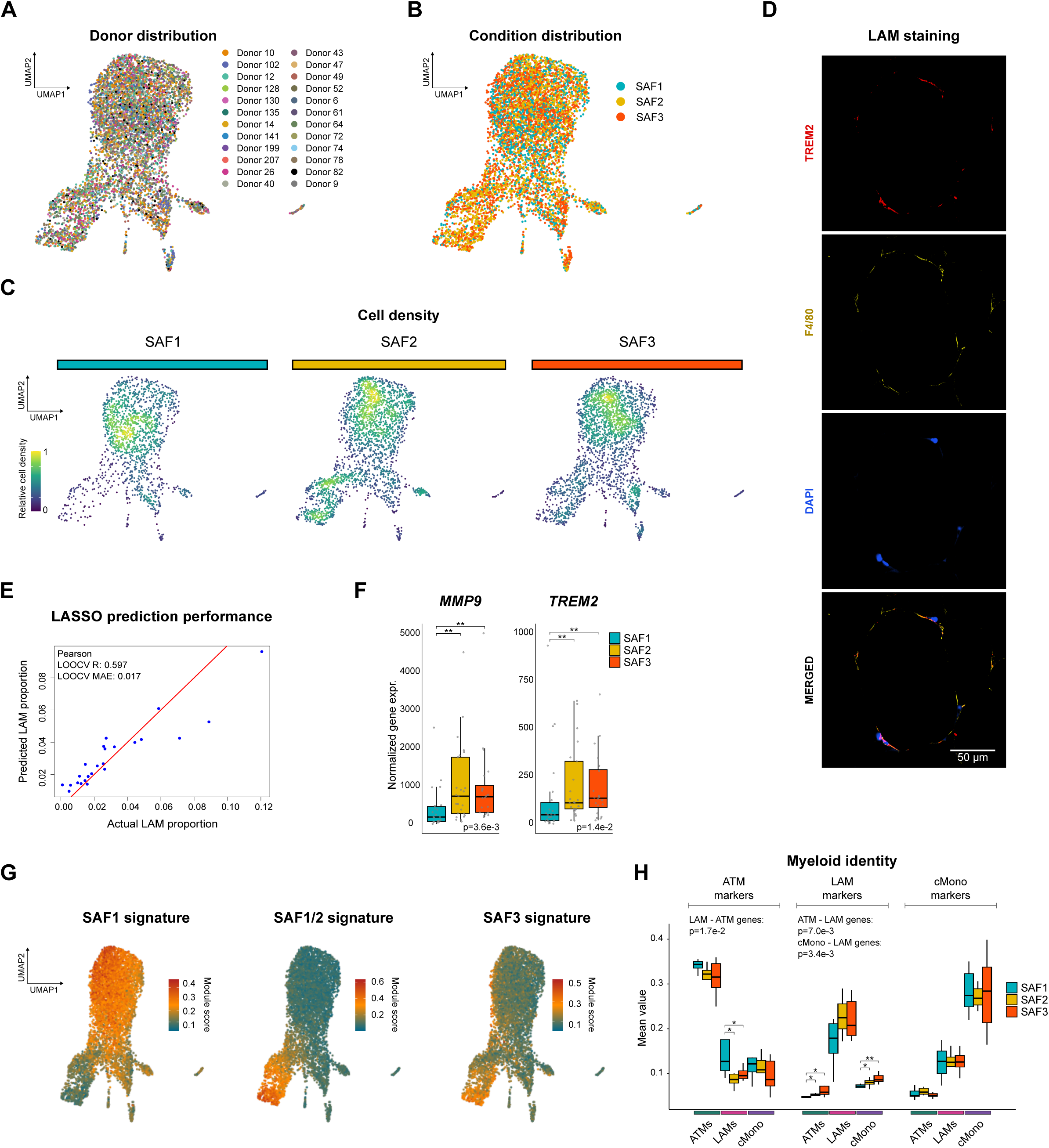
| Lipid-associated macrophages are activated in MASLD. **A–B**, UMAP visualization of myeloid immune cell nuclei from all donors, colored by individual donor (**A**) and SAF score (**B**). **C**, Density plot showing distribution of myeloid subpopulations across SAF categories. **D**, Immunofluorescent images showing overlap between TREM2 (red), F4/80 (yellow) and DAPI (blue) in SAT. **E**, Scatter plot comparing known cell type proportions from matched snRNA-seq data (x-axis) with predicted proportions from bulk RNA-seq data (y-axis) for LAMs, estimated using a LASSO regression model. Red line indicates regression fit. **F**, Normalized expression of selected LAM marker genes from bulk RNA-seq data of 67 female individuals stratified by SAF score (all: S1, n=22; S2, n=25, S3, n=20). **G,** UMAP visualization of UCell module scores in myeloid subpopulations for different myeloid gene clusters (as defined in Figure 2A). **H**, Normalized pseudobulk expression of markers for ATM, LAMs, and classical monocytes across SAF categories in indicated myeloid subpopulations. Boxplot data: center line, median; box limits, upper and lower quartiles; whiskers, 1.5x IQR. A KW test, followed by a post-hoc Wilcoxon rank-sum test with BH correction for multiple testing, was performed to assess significance between SAF diagnosis for **F** and **H** *, p<0.05; **, p<0.01). Model performance was evaluated using leave-one-out cross-validation (LOOCV), and accuracy was quantified by Pearson correlation and mean absolute error (MAE) in **E**. ATM, adipose tissue-resident macrophages; cMono, classical monocytes; LAMs, lipid-associated macrophages.

**Figure S4.**
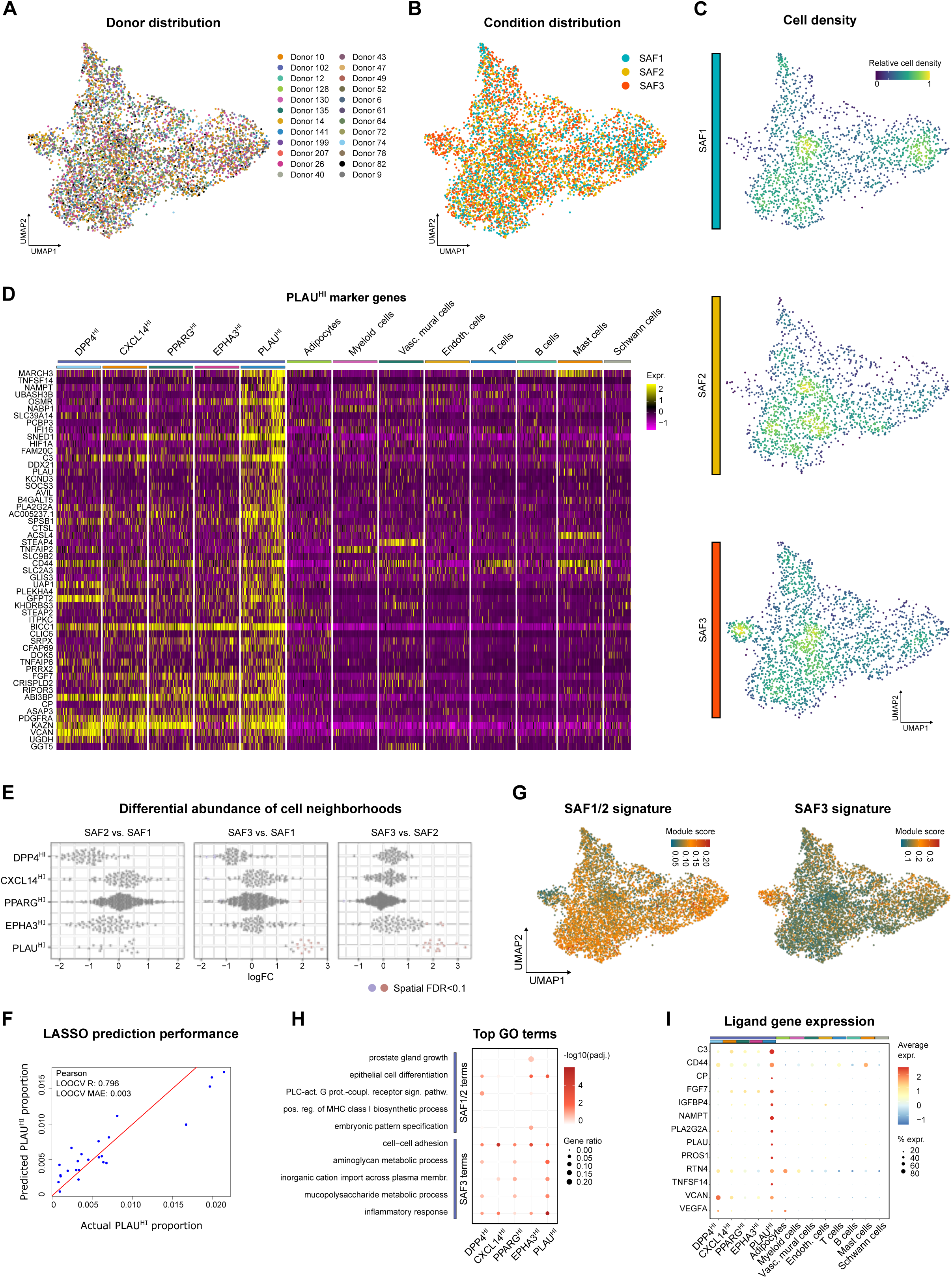
| ASPC remodeling in SAT with increasing MASLD severity. **A–B**, UMAP visualization of ASPC nuclei from all donors, colored by individual donor (**A**) and SAF score (**B**). **C**, Density plot showing distribution of ASPC subpopulations. **D**, Heatmap of normalized expression of tissue-wide *PLAU*^HI^ ASPC marker genes. **E**, Distribution of ASPC subpopulation neighborhoods for indicated SAF score comparisons. Blue, downregulated; Red, upregulated; Grey, unregulated. **F**, Scatter plot comparing known *PLAU*^HI^ ASPC proportions from matched snRNA-seq data (x-axis) with predicted proportions from bulk RNA-seq data (y-axis), estimated using a LASSO regression model. Red line indicates regression fit. **G**, UMAP visualization of UCell module scores in ASPC subpopulations for different ASPC gene clusters (as defined in Figure 2B). **H**, Enrichment of genes linked to top GO biological process pathways (as defined in Figure 2E) within ASPC subpopulations. **I**, Normalized expression of top predicted *PLAU*^HI^ ASPC ligands in indicated ASPC subpopulations. Model performance was evaluated using LOOCV, and accuracy was quantified by Pearson correlation and MAE in **F**. ASPCs, adipogenic stromal and progenitor cells.

**Figure S5.**
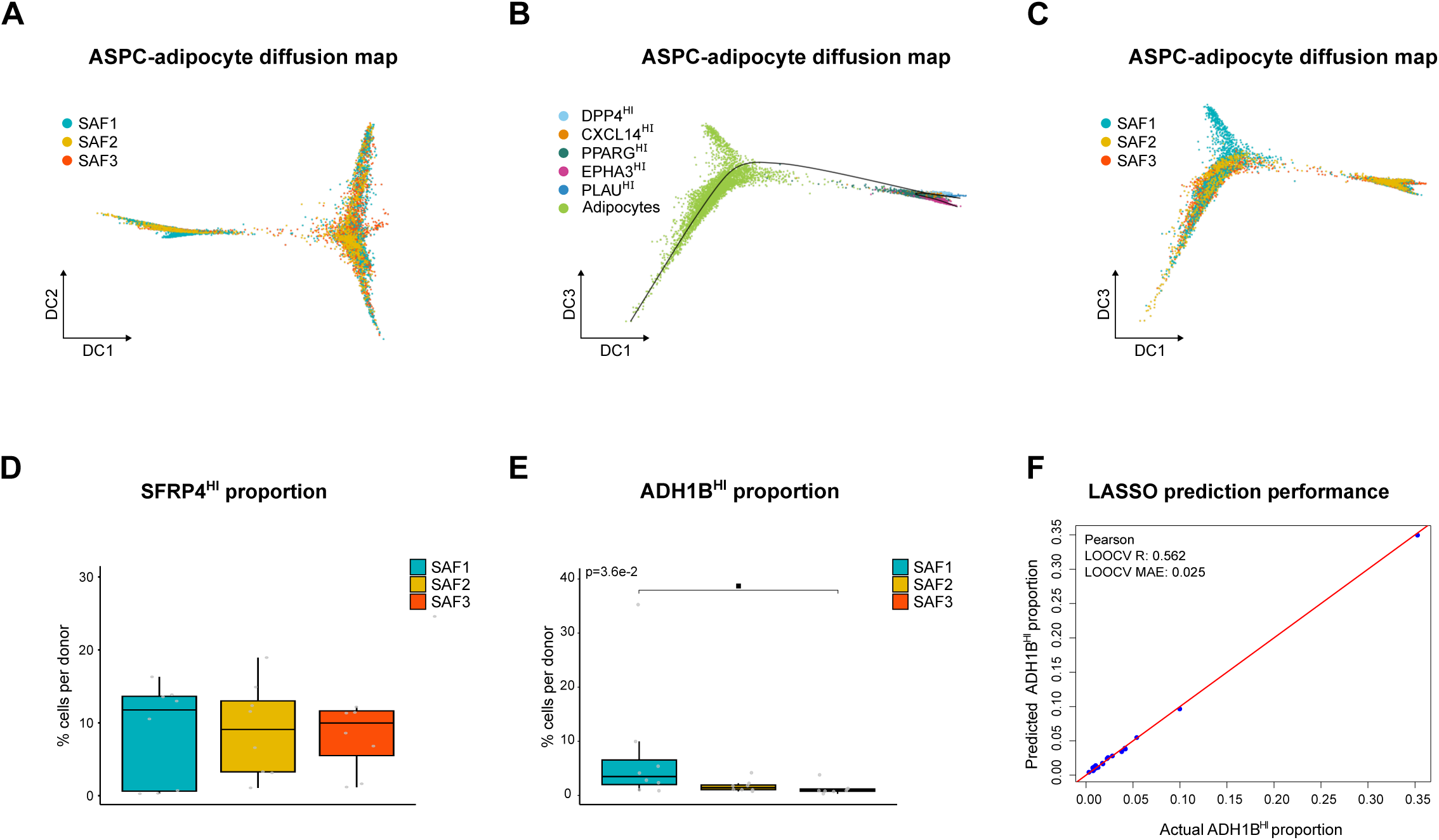
| *ADH1B*^HI^ adipocytes are lost in individuals with MASLD. **A**, Diffusion map (DC1 vs DC2) of ASPC and adipocyte nuclei from all donors, colored by SAF score. **B**, Diffusion map (DC1 vs DC3) of ASPC and adipocyte nuclei from all donors. **C**, Diffusion map (DC1 vs DC3) of ASPC and adipocyte nuclei from all donors, colored by SAF score. **D**, Proportional abundance of *SFRP4*^HI^ (**D**) and *ADH1B*^HI^ adipocytes (**E**) across SAF categories in the human abdominal SAT dataset (n=8 per condition). **f**, Scatter plot comparing known proportions of *ADH1B*^HI^ adipocytes from matched snRNA-seq data (x-axis) with predicted proportions from bulk RNA-seq data (y-axis), estimated using a LASSO regression model. Red line indicates regression fit. Boxplot data: center line, median; box limits, upper and lower quartiles; whiskers, 1.5x IQR. A KW test, followed by a post-hoc Wilcoxon rank-sum test with BH correction for multiple testing, was performed to assess significance between SAF diagnosis for **D-E** (▪, p<0.1). Model performance was evaluated using LOOCV, and accuracy was quantified by Pearson correlation and MAE in **F**. ASPCs, adipogenic stromal and progenitor cells.

**Figure S6.**
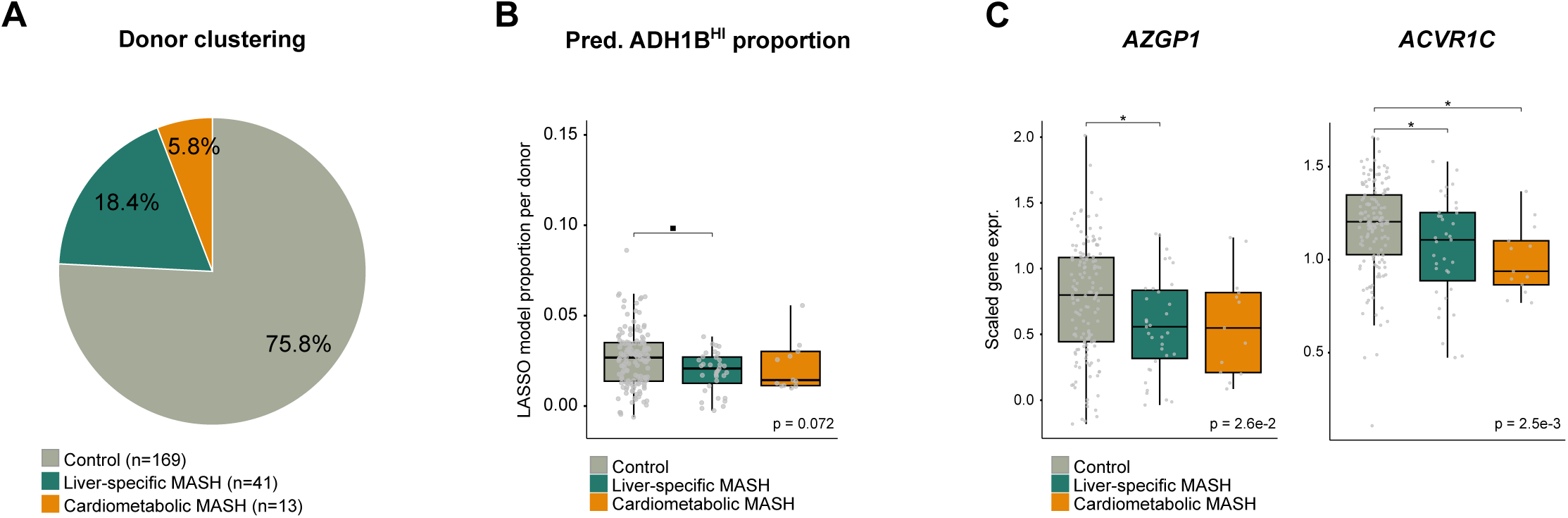
| Adipose tissue subpopulations associate differently with MASH endotypes. **A**, Pie chart showing the percentage of individuals in control and MASH endotype clusters for the expanded cohort (Control, n=169; LS, n=41, CM, n=13). **B**, LASSO-predicted *ADH1B*^HI^ adipocyte proportions in bulk RNA-seq data in control and MASH endotype clusters (Control, n=146; LS, n=36, CM, n=13). **C**, Normalized expression of selected *ADH1B*^HI^ adipocyte marker genes from bulk RNA-seq data in control and MASH endotype clusters (Control, n=146; LS, n=36, CM, n=13). Boxplot data: center line, median; box limits, upper and lower quartiles; whiskers, 1.5x IQR. A KW test, followed by a post-hoc Wilcoxon rank-sum test with BH correction for multiple testing, was performed to assess significance between SAF diagnosis for **B-C** (▪, p<0.1; *, p<0.05).

